# The effects of GC-biased gene conversion on patterns of genetic diversity among and across butterfly genomes

**DOI:** 10.1101/2020.11.10.376566

**Authors:** Jesper Boman, Carina F. Mugal, Niclas Backström

## Abstract

Recombination reshuffles the alleles of a population through crossover and gene conversion. These mechanisms have considerable consequences on the evolution and maintenance of genetic diversity. Crossover, for example, can increase genetic diversity by breaking the linkage between selected and nearby neutral variants. Bias in favor of G or C alleles during gene conversion may instead promote the fixation of one allele over the other, thus decreasing diversity. Mutation bias from G or C to A and T opposes GC-biased gene conversion (gBGC). Less recognized is that these two processes may –when balanced– promote genetic diversity. Here we investigate how gBGC and mutation bias shape genetic diversity patterns in wood white butterflies (*Leptidea* sp.). This constitutes the first in-depth investigation of gBGC in butterflies. Using 60 re-sequenced genomes from six populations of three species, we find substantial variation in the strength of gBGC across lineages. When modeling the balance of gBGC and mutation bias and comparing analytical results with empirical data, we reject gBGC as the main determinant of genetic diversity in these butterfly species. As alternatives, we consider linked selection and GC content. We find evidence that high values of both reduce diversity. We also show that the joint effects of gBGC and mutation bias can give rise to a diversity pattern which resembles the signature of linked selection. Consequently, gBGC should be considered when interpreting the effects of linked selection on levels of genetic diversity.

## Introduction

The neutral theory of molecular evolution postulates that the majority of genetic differences within and between species are due to selectively neutral variants (Kimura 1983; Jensen, et al. 2019). Consequently, the level of genetic variation within populations (θ) is expected to predominantly be determined by the effective population size (*N*_*e*_) and the mutation rate (μ) according to the following relationship: θ = 4*N*_*e*_μ. Indeed, differences in life-history characteristics (as a proxy for *N*_*e*_) have been invoked as explanations for the interspecific variation in genetic diversity among animals (Romiguier, et al. 2014). Also among butterflies, body size is negatively associated with genetic diversity (Mackintosh, et al. 2019). Usually the population size estimated from genetic diversity measures is lower than expected based on the classic neutral model and census population size, *N*_*c*_ (Lewontin 1974; Kimura 1983; Nevo, et al. 1984; Frankham 1995). This observation has been called Lewontin’s paradox (*N*_*e*_ < *N*_*c*_) and may be caused by more efficient selection and subsequently reduced genetic diversity in large compared to small populations (Corbett-Detig, et al. 2015). In particular, selection affects the allele frequency of linked neutral sites (commonly referred to as linked selection or genetic draft) and reduces their diversity (Maynard Smith and Haigh 1974; Charlesworth, et al. 1993).

However, linked selection in itself is not necessarily the solution to Lewontin’s paradox. It has been noted that *N*_*e*_ = *N*_*c*_ is true only for a population in mutation-drift equilibrium (Galtier and Rousselle 2020). Furthermore, changes in population size may amplify the effects of linked selection and the relative importance of selection and demography is an ongoing debate (Corbett-Detig, et al. 2015; Coop 2016; Kern and Hahn 2018; Jensen, et al. 2019). This debate concerns the fate and forces affecting an allele while segregating in a population. While this is important for resolving Lewontin’s paradox, it only addresses variation in *N*_*e*_, which is but a part of the puzzle of genetic diversity. As noted above, variation in the occurrence of mutations also influences genetic diversity. The general pattern observed is a negative relationship between mutation rate and *N*_*e*_ (Lynch, et al. 2016). This is explained by the observation that the distribution of fitness effects of new mutations are dominated by deleterious mutations which leads to a selective pressure for reducing the overall mutation rate (Eyre-Walker and Keightley 2007; Lynch, et al. 2016). However, mutation rates vary only over roughly one order of magnitude in multicellular eukaryotes (Lynch, et al. 2016) and appear less important than *N*_*e*_ for interspecific differences in genetic diversity.

Genetic diversity can also vary among genomic regions. The determinants of such regional variation are currently debated, but variation in mutation rate (Hodgkinson and Eyre-Walker 2011; Smith, et al. 2018) and linked selection have both been considered (Cutter and Payseur 2013; Corbett-Detig, et al. 2015). Higher rates of recombination are expected to reduce the decline in diversity experienced by sites in the vicinity of a selected locus. Begun and Aquadro (1992) showed for example that genetic diversity was positively correlated with the rate of recombination in *Drosophila melanogaster.* Their finding validated the impact of selection on linked sites, previously predicted by theoretical work (reviewed in Comeron 2017). Since then, multiple studies have found a positive association between recombination rate and genetic diversity (e.g. Begun and Aquadro 1992; Nachman 1997; Kraft, et al. 1998; Cutter and Payseur 2003; Stevison and Noor 2010; Lohmueller, et al. 2011; Rao, et al. 2011; Langley, et al. 2012; Cutter and Payseur 2013; Mugal, et al. 2013; Burri, et al. 2015; Corbett-Detig, et al. 2015; Wallberg, et al. 2015; Martin, et al. 2016; Pouyet, et al. 2018; Castellano, et al. 2019; Rettelbach, et al. 2019; Talla, Soler, et al. 2019). The positive correlation between diversity and recombination may, however, be caused by factors other than selection on linked sites. Recombination may for instance be mediated towards regions of higher genetic diversity (Cutter and Payseur 2013), or have a direct mutagenic effect (Hellmann, et al. 2005; Arbeithuber, et al. 2015; Halldorsson, et al. 2019). Additionally, analytical evidence suggests that the interplay between mutation bias and a recombination-associated process, GC-biased gene conversion (gBGC), can increase nucleotide diversity (McVean and Charlesworth 1999). GC-biased gene conversion in itself will like directional selection reduce diversity of segregating variants. If we additionally consider the long-term effect of gBGC and the concomitant increase in GC content, then genetic diversity may rise as a consequence of gBGC through increased mutational opportunity in the presence of an opposing mutation bias (McVean and Charlesworth 1999). To fully understand the effects of recombination on genetic diversity we must therefore consider both gBGC and opposing mutation bias, in addition to the much more recognized influence of linked selection. In other words, what relationship do we expect between recombination and genetic diversity in the presence of non-adaptive forces such as gBGC and mutation bias?

To understand the mechanistic origins of gBGC we must first consider gene conversion, a process arising from homology directed DNA repair during recombination. Gene conversion is the unilateral exchange of genetic material from a donor to an acceptor sequence (Chen, et al. 2007). A recombination event is initiated by a double-strand break which is repaired by the cellular machinery using the homologous chromosome as template sequence. If there is a sequence mismatch within the recombination tract, gene conversion may occur (Chen, et al. 2007). Mismatches in heteroduplex DNA are repaired by the mismatch-repair machinery (Chen *et al.* 2007). Importantly, G/C (strong, S, three-hydrogen bonds) to A/T (weak, W, two hydrogen bonds) mismatches can have a resolution bias in favor of S alleles resulting in gBGC, a process that can alter base composition and genetic diversity (Nagylaki 1983a, b; Marais 2003; Duret and Galtier 2009; Mugal, et al. 2015). Direct observations of gBGC are restricted to a small number of taxa, such as human (Arbeithuber, et al. 2015), baker’s yeast (*Saccharomyces cerevisiae*) (Mancera, et al. 2008), collared flycatcher (Smeds, et al. 2016) and honey bees (Kawakami, et al. 2019). Indirect evidence exists for a wider set of species, including arthropods such as brine shrimp (*Artemia franciscana*) and butterflies from the Hesperidae, Pieridae and Nymphalidae families (Eyre-Walker 1999; Perry and Ashworth 1999; Meunier and Duret 2004; Spencer, et al. 2006; Muyle, et al. 2011; Pessia, et al. 2012; Glémin, et al. 2015; Galtier, et al. 2018).

The strength of gBGC can be measured by the population-scaled parameter *B* = 4*N*_*e*_*b*, where *b* = *ncr* is the conversion bias, which is dependent on the average length of the conversion tract (*n*), the transmission bias (*c*), and the recombination rate per site per generation (*r*) (Glémin, et al. 2015). This means that we can expect a stronger impact of gBGC in larger populations and in genomic regions of high recombination. Nagylaki (1983a) showed that we can understand gBGC in terms of directional selection, i.e. the promotion of one allele over another. This leads to a characteristic derived allele frequency (DAF) spectrum, in which an excess of W→S alleles- and a concomitant lack of S→W alleles, are segregating at high frequencies in the population. Nevertheless, the overall number of S→W polymorphism is expected to be higher in most species because of the widely observed S→W mutation bias, partially caused by the hypermutability of methylated cytosines in the 5’-CpG-3’ dinucleotide context (Lynch 2007). Preventing the fixation of ubiquitous and possibly deleterious S→W mutations have been proposed as one of the ultimate causes for gBGC (Brown and Jiricny 1987; Birdsell 2002; Duret and Galtier 2009). However, while gBGC reduces the mutational load it may also confer a substitutional load by favoring deleterious W→S alleles (Duret and Galtier 2009; Glémin 2010; Mugal, et al. 2015). This effect has led some authors to describe gBGC as an “Achilles heel” of the genome (Duret and Galtier 2009; Mugal, et al. 2015). Detailed analysis of a larger set of taxonomic groups is needed to understand the prevalence and impact of gBGC. There is also limited knowledge about the variation in the strength of gBGC within and between closely related species (Borges, et al. 2019).

Here, we investigate the dynamics of gBGC in butterflies and characterize the effect of gBGC on genetic diversity. We use whole-genome re-sequencing data from 60 individuals from six populations of three species of wood whites (genus *Leptidea*). Wood whites show distinct karyotype- and demographic differences both within and among species (Dincă, et al. 2011; Lukhtanov, et al. 2011; Dincă, et al. 2013; Lukhtanov, et al. 2018; Talla, Johansson, et al. 2019; Talla, Soler, et al. 2019). This includes, *L. sinapis,* which has the greatest intraspecific variation in diploid chromosome number of any animal, from 2n = 57,58 in southeastern Sweden to 2n =106-108 in northeastern Spain (Lukhtanov, et al. 2018). Our objectives are threefold. First, we infer the strength and determinants of gBGC variation among *Leptidea* populations. Second, we investigate the patterns of gBGC and mutation bias across the genome, its determinants and association with GC content. Third, we detail the effect of gBGC and opposing mutation bias on genetic diversity across a GC gradient and consider the impact of linked selection and GC content itself as determinants of genetic diversity.

## Materials and methods

### Samples, genome and population resequencing data

The samples and population resequencing data used in this study were originally presented in Talla, et al. (2017). In brief, 60 male *Leptidea* butterflies from three species and six populations were analyzed. For *L. sinapis* (Figure 1B), 30 individuals were sampled: 10 from Kazakhstan (Kaz-sin), 10 from Sweden (Swe-sin) and 10 from Spain (Spa-sin). 10 *L. reali* were sampled in Spain (Spa-rea) and 10 *L. juvernica* per population were collected in Ireland (Ire-juv) and Kazakhastan (Kaz-juv), respectively (Figure 1A). Reads from all 60 sampled individuals were mapped to a previously available genome assembly of an inbred, male, Swedish *L. sinapis* (scaffold N50 = 857 kb) (Talla, et al. 2017). Detailed information on SNP calling can be found in Talla, Johansson, et al. (2019). Chromosome numbers for each population (if available) or species were obtained from the literature (Dincă, et al. 2011; Lukhtanov, et al. 2011; Šíchová, et al. 2015; Lukhtanov, et al. 2018).

**Figure 1.**
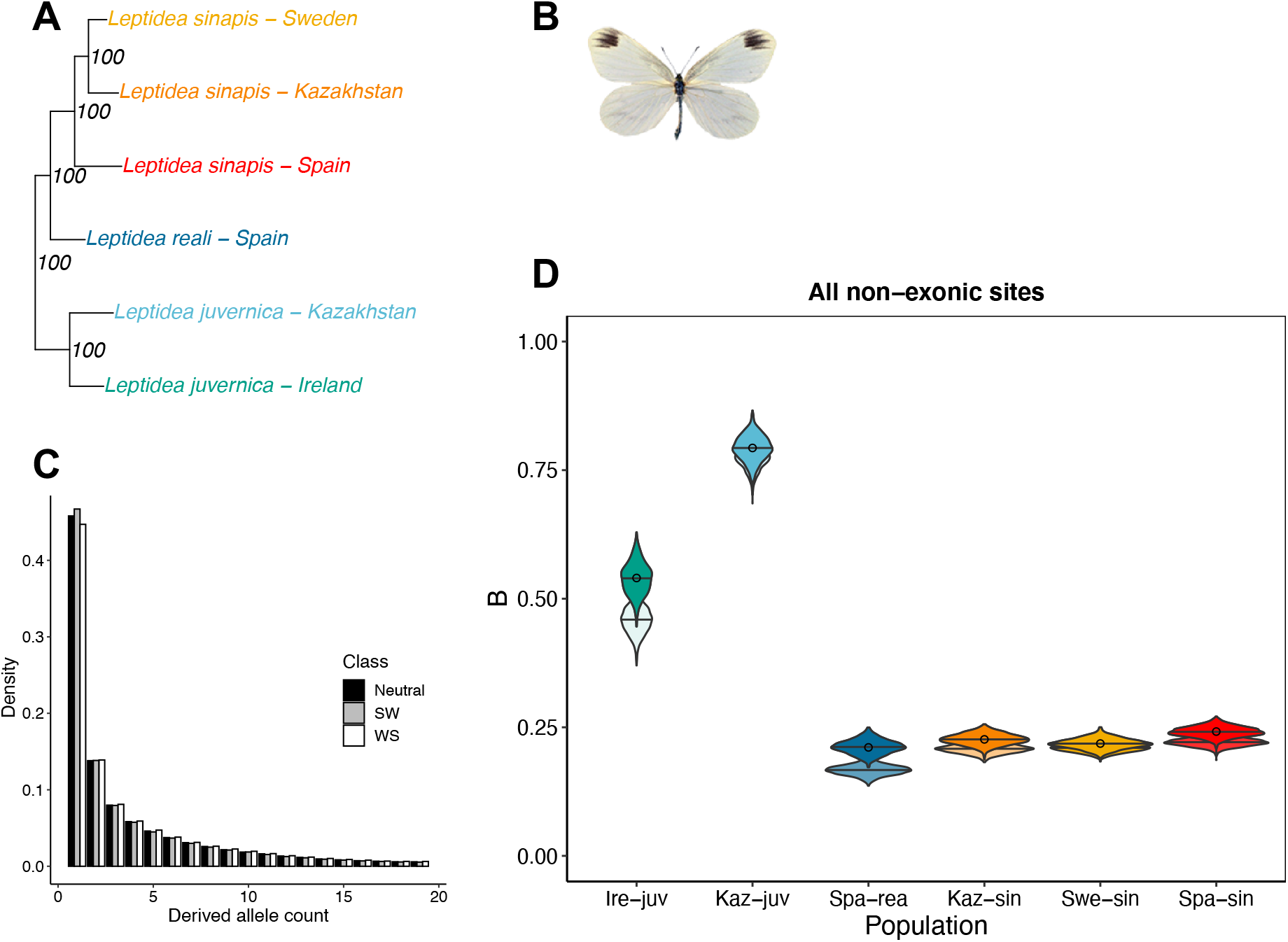
*Leptidea* butterflies show variation in the genome-wide strength of gBGC. A) Phylogeny of the six *Leptidea* populations included in this study. Node values represent support from 100 bootstrap replicates on sites. The phylogeny in A) is based on a subtree from a maximum-likelihood phylogeny used as a starting tree in Figure 1 of Talla *et al.* (2017). B) Mounted specimen of *Leptidea sinapis*. C) DAF spectra for polarized non-exonic SNPs of the Swedish *L. sinapis* population split in categories S→W (SW), W→S (WS) and GC-neutral. D) Estimates of the population-scaled coefficient of gBGC (*B =* 4*N*_*e*_*b*). Circles represent point estimates from the original DAF spectra using model M1*, bars are mean values of *B* for the 1,000 bootstrap replicates of sites. Overlain and opaque violins are bootstrapped values for model M1* and underlain, transparent violins are estimates for model M1.

### Filtering and polarization of SNPs

Allele counts for each population were obtained using *VCFtools* v. 0.1.15 (Danecek, et al. 2011). Only non-exonic, biallelic SNPs with no missing data for any individual, and in regions not masked by *RepeatMasker* in the *L. sinapis* reference assembly (Talla, et al. 2017; Talla, Johansson, et al. 2019), were kept for downstream. The rationale behind excluding exonic SNPs was to minimize the impact of selection on the allele frequencies, and SNPs in repetitive regions were excluded because of the reduced ability for unique read mapping (Sexton and Han 2019), and their higher potential for ectopic gene conversion, which deserve a separate treatment (Roy, et al. 2000; Chen, et al. 2007). Sex-chromosome linked SNPs were considered like any other SNP. The lack of recombination in female meiosis in butterflies (Maeda 1939; Suomalainen, et al. 1973; Turner and Sheppard 1975) and the reduced effective population size (*N*_*e*_, three Z chromosomes per four autosomes [*A*]) cancel out (Charlesworth 2012). This leaves only their relative recombination rate (*r*) affecting intensity of gBGC (*B*), assuming that effective sex ratios are equal and that conversion tract length (*n*) and transmission bias (*c*) are equal between Z and A.

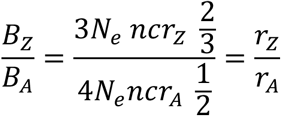

SNPs were polarized using invariant sites in one or two outgroup populations, again allowing no missing data (Table S1). We denote this polarization scheme “strict”. We also tested a more “liberal” polarization approach where only the individual with highest average read depth per outgroup population was used to polarize SNPs, allowing for one missing allele per individual. Mean read depth per individual was obtained using *VCFtools* v. 0.1.15 (Danecek, et al. 2011). The liberal polarization scheme was mainly used to test the impact of polarization on estimation of the mutation bias (λ) of S→W mutations over W→S mutations given mutational opportunity (Table S1). The strict polarization was used for all analysis unless otherwise stated. We considered alternative (i.e. not in the reference genome) alleles as the ancestral allele if all outgroup individual(s) were homozygous for that allele (strict polarization and liberal polarization).

Derived allele frequency spectra of segregating variants were computed for the following categories of mutations; GC-conservative/neutral (S→S and W→W, here denoted N→N), strong to weak (S→W) and weak to strong (W→S). All alternative alleles inferred as ancestral alleles were used to replace the inferred derived reference allele to make a model of an ancestral genome using *BEDTools* v. 2.27.1 *maskfasta* (Quinlan & Hall 2010). This method leverages the information from invariant sites in all sequenced individuals to decrease the reference bias when calculating GC content. However, the ancestral genome was biased towards *L. sinapis* given that it both served as a reference genome and had more polarizable SNPs than the *L. reali* and *L. juvernica* populations (Table S1).

### Inferring GC-biased gene conversion from the DAF spectrum

To estimate the strength of gBGC, we utilized a population genetic maximum likelihood model (Muyle, et al. 2011; Glémin, et al. 2015), implemented as a notebook in *Mathematica v. 12.0* (Wolfram Research 2019). The model jointly estimates the S→W mutation bias (λ) and the population-scaled coefficient of gBGC (*B* = 4*N*_*e*_*b*), in which *b* is the conversion bias. To account for demography, the model introduces a nuisance parameter (*r*_*i*_) per derived allele frequency class (*i*) following Eyre-Walker, et al. (2006). The model also estimates the genetic diversity of N→N and W→S spectra (θ_N_ and θ_WS_ respectively) and computes an estimate of the skewness of S→W and W→S alleles in the folded site frequency spectrum. We applied four of the implemented models, i.e. M0, M0*, M1 and M1*, as the more extended models have large variance without prior information on heterogeneity of recombination intensity at a fine scale (Glémin, et al. 2015), which is currently lacking for Lepidoptera. The M0 model is a null model that evaluates the likelihood of the observed DAF spectrum for a population genetic model without gBGC (i.e. *B* = 0). M1 extends this model by including gBGC via the parameter *B*. M0* and M1* are extensions of M0 and M1, respectively, where one additional parameter per mutation class is incorporated, to account for polarization errors. We analyzed separately all non-exonic sites, and excluding- or including ancestral CpG-prone sites, meaning trinucleotides including the following dinucleotides: CG, TG, CA, NG, TN, CN, NA centered on the polarized variant. N here means either a masked or unknown base. Following Glémin, et al. (2015), we used GC content as a fixed parameter in the maximum likelihood estimation. GC content in the repeat- and gene-masked ancestral genome model was determined by the *nuc* program in the *BEDTools* v.2.27.1 suite. Coordinates of repeats and exons (including introns and UTR regions if available) were obtained from Talla, et al (2017) and Leal, et al (2018), respectively. The number of G and C bases at ancestral CpG-prone sites were computed using a custom script and subtracted from the GC of all non-exonic sites to obtain the GC content for the set excluding ancestral CpG-prone sites.

### GC centiles

The polarized non-repetitive, non-exonic SNPs of each population were divided into 100 ranked bins based on local GC content (GC centiles) in the repeat- and exon-masked ancestral genome. This means, all GC centiles represented unequally sized chunks of the genome with equal numbers of polarizable SNPs. The GC content was estimated in 1 kb windows of the reconstructed and repeat- and exon-masked ancestral genome (described above) using *BEDTools* v. 2.27.1 *nuc* (Quinlan and Hall 2010), correcting for the number of N bases. To calculate the overall GC content of a centile, we summed the GC content of each 1 kb window. Separate DAFs were created per centile and parameters of gBGC and mutation bias were estimated with the models previously described. We also estimated the genetic diversity per GC centile and population using the average pairwise differences (nucleotide diversity, π), and excluded masked bases when averaging. We calculated π for all sites without any missing data, separately for each population, using 1 as value for the max missing (-mm) parameter in the *-- site-pi* function of *VCFtools* v. 0.1.15 (Danecek, et al. 2011). We also calculated separate π for polarized sites belonging to the following mutation categories (S→S), (W→W), (S→W) and (W→S) for each population and centile, using a custom function in *R* (R Core Team 2020). To average π, we used the number of unmasked bases within the range of GC values defined by each centile. The proportion of coding bases (CDS density) was used as a proxy for the impact of linked selection in general, and background selection in particular. CDS density was estimated separately for each population and centile by aggregating the CDS content across all 1 kb windows for a particular centile. A custom-made script was used to assess the impact of read depth on the pattern of π across GC centiles (Figure 4D). This script combined *BEDTools* v. 2.27.1 (Quinlan and Hall 2010) *complement, genomecov* and *intersect* to calculate the read depth per non-masked base pair. Average read-depth per individual and centile was then plotted against GC content to qualitatively assess if the population specific patterns followed what was observed for the association between π and GC.

### Model of the effect of gBGC and mutation bias on genetic diversity

We considered a model in which the effect of gBGC (*B*) and mutation bias (λ) determines the level of π relative to a reference case where *B* = 0 (McVean and Charlesworth 1999),

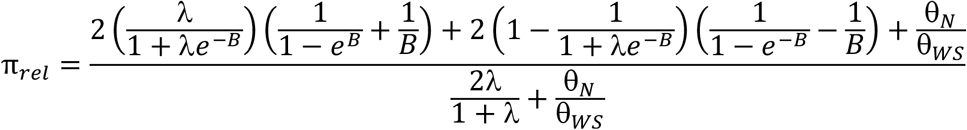

The numerator consists of three terms each describing the relative contributions of S→W, W→S and N→N mutations. GC-changing mutations have a diversity determined by λ and *B* while the contribution of GC-conservative/neutral mutations are affected by the ratio of N→N diversity (θ_N_) over W→S diversity (θ_WS_). The denominator achieves the standardization for the reference case (see above). The model assumes gBGC-mutation-drift (GMD) equilibrium. From an empirical perspective this means that π_rel_ is the predicted π relative to the reference case (*B* = 0) when the observed GC content is at a value determined by gBGC and mutation bias (1/1+λe^−*B*^). Fitting the GMD model relies on obtaining a neutral reference π value unaffected by demographic fluctuations in population size, selection or gBGC. Such a value is unattainable, except for the most well-studied model organisms (Pouyet, et al. 2018). Maximum observed genetic diversity, π_max_, could be used as a proxy for neutral diversity which should be reasonable if all centiles are reduced below their neutral value through linked selection (Torres, et al. 2020). Another approach, which we employ here, is to fit the model without estimating a neutral reference π. This allows us to estimate how *B, λ* and the relative amount of GC-changing mutations affect π_*rel*_.

### Statistical analyses

All statistical analyses were performed using *R* v. 3.5.0-4.0.2 (R Core Team 2020). Linear models and correlations were performed using default packages in R. Phylogenetic independent contrasts (Felsenstein 1985) were performed using the *pic()* function in the package *ape* (Paradis and Schliep 2018). This package was also used to depict the phylogeny in Figure 1A. Other plots were either made using base *R* or the *ggplot2* package (Wickham 2016).

## Results

### Patterns of gBGC among populations and species

To infer the strength of gBGC in the different *Leptidea* populations (Figure 1A, B), separate DAFs for segregating non-exonic variants for each category of mutations (N→N, S→W and W→S) were calculated (see example from Swe-sin in Figure 1C). We used the four basic population genetic models developed by Glemin et al. (2015) to obtain maximum likelihood estimates of the intensity of gBGC (*B* = 4N_e_b). The GC content in the ancestral genome was ~0.32. For all populations, the M1 model had a better fit than the M0 model (likelihood-ratio tests (LRT) upper-tailed χ^2^; α = 0.05; df = 1), which indicates that gBGC is a significant evolutionary force in *Leptidea* butterflies (Figure 1D). The quantitative results from the M1 and M1* models were overall congruent, and M1* had a better fit for all populations except Swe-sin (LRT upper-tailed χ^2^; α = 0.05; df = 3). When taking all non-exonic sites into consideration and applying model M1*, Spa-rea and Swe-sin had the lowest *B* (0.21), followed by Kaz-sin (*B* = 0.22). Spa-sin, the population with the largest number of chromosomes (Figure 2B), had a marginally higher *B* (0.24). All these estimates were lower than Irish-(Ire-juv) and Kazakhstani (Kaz-juv) *L. juvernica* with *B* = 0.54 and *B* = 0.79, respectively (Table S2).

**Figure 2.**
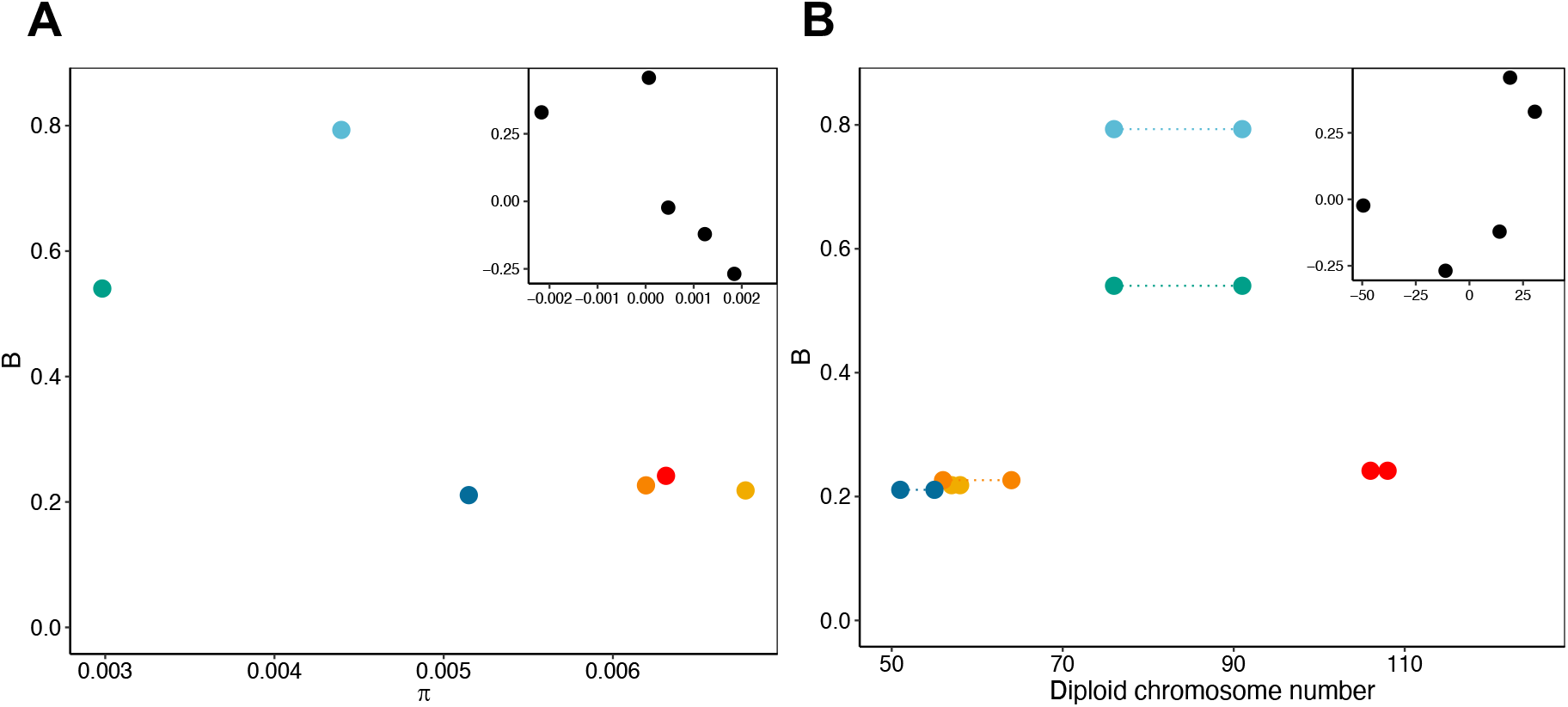
Determinants of variation in the strength of gBGC among populations. A) Relationship between π and *B*. B) Relationship between diploid chromosome number and *B*. Points in B) show lowest and highest estimate of diploid chromosome number for each population. Colors represent the populations shown in Figure 1. Insets in A) and B) show phylogenetically independent contrasts of each respective axis variable based on the phylogeny in Figure 1A. Contrasts for diploid chromosome number were based on midpoint value.

We tried an alternative more “liberal” polarization (only 2 outgroup individuals, see *Materials and Methods*) to test the impact of the polarization scheme on the estimates from the gBGC model. The results were qualitatively similar but the polarization error rates were inflated compared to the “stricter” polarization scheme (Table S2, Text S1). Thus, we used the “strict” polarization scheme for subsequent analyses unless otherwise stated. We also tested the impact of including and excluding ancestral CpG-prone sites as they may impact the estimation of the S→W mutation bias (λ) and *B* (Text S1). All populations except Kaz-juv had the highest estimate of λ at ancestral CpG-prone sites, followed by all non-exonic sites and lowest when excluding ancestral CpG sites (Table S2). This difference could be caused by hyper-mutagenic methylated cytosines but the level of DNA methylation observed in Lepidopteran taxa is low (Bewick, et al. 2017; Jones, et al. 2018; Provataris, et al. 2018).

### Determinants of gBGC intensity variation among populations and species

The strength of gBGC is dependent on *N*_*e*_ and the conversion bias *b* = *ncr*. Given that transmission bias, *c,* and conversion tract length, *n*, require sequencing of pedigrees, we here focus on variation in genome-wide recombination rate, *r* to assess variation in *b.* To understand the relative importance of *N*_*e*_ and *r*, we correlated *B* with π (as a proxy for *N*_*e*_) and diploid chromosome number (as a proxy for genome-wide recombination rate). Neither genetic diversity, (π*; p* ≈ 0.13, adjusted *R*^*2*^ ≈ 0.45), nor diploid chromosome number (*p* ≈ 0.35, *R*^*2*^ ≈ 0.05), significantly predicted variation in *B* among species in phylogenetically independent contrasts (Figure 2C, D). Since Spanish *L. sinapis* likely experienced massive chromosomal fission events recently (Lukhtanov, et al. 2011; Talla, Johansson, et al. 2019; Lukhtanov, et al. 2020), it is possible that *B* is below its equilibrium value in this population. Excluding Spa-sin yielded a marginally significant positive relationship between chromosome number and the intensity of gBGC (*p* ≈ 0.07, *R*^*2*^ ≈ 0.79).

### Level of mutation bias varies among *Leptidea* species

The GC content is determined by the relative fixation of S→W and W→S mutations (Sueoka 1962), which is governed by the balance of a mutation bias from S→W over W→S, and a fixation bias from W→S over S→W. The latter may be caused by gBGC only, but may also be observed at synonymous sites due to selection for preferred codons (Duret and Mouchiroud 1999; Clément, et al. 2017; Galtier, et al. 2018). Protein coding genes make up only 3.7 % of the *L. sinapis* genome (Talla, Soler, et al. 2019) and potential selection on codon usage will hence only affect genome-wide base composition marginally in this species. Using the DAF spectra of different mutation classes allows not only estimation of *B,* but also the mutation bias, λ (Muyle, et al. 2011; Glémin, et al. 2015). We found that λ (estimated from model M1*) varied from 2.94 (e.g. Spa-sin) to 4.09 (Kaz-juv) (Table S2). Applying the M1 model gave similar results. It is possible that the polarization scheme which only allowed private alleles for the *L. juvernica* populations, contributes to their high value of λ. To test this, we polarized genome-wide, non-exonic SNPs using the individuals from the other two species with the highest read depth as outgroups. The resulting λ were ~3.5 and ~3 for Kaz-juv and Ire-juv respectively and ~3 for the *L. reali* and *L. sinapis* populations, with only minor differences in λ between the M1 and M1* models for all populations (Table S2). This indicates that the strict polarization scheme shapes the DAF spectrum in a way unaccountable for by the demographic *r*_*i*_ parameters of the model. However, the polarization scheme alone cannot explain the higher λ observed in Kaz-juv compared to the other populations (see Text S1 for further discussion).

### Patterns and determinants of gBGC and GC content across the genome

To understand the effects of gBGC throughout the genome, we partitioned the polarized SNPs into centiles based on their local (1kb) GC content in the ancestral genome. The number of SNPs in each centile ranged from 2,661 in Ire-juv to 21,140 in Spa-sin (Table S1). The models were compared using LRTs on the average difference of all centiles between the reduced (M0) and full (M1) model and between the models excluding (M1) or including (M1*) polarization error parameters. M0 could not be rejected in favor of M1 for both Ire-juv and Spa-rea. It is possible that the lower number of SNPs per GC centile in these populations increases variance and thus reduces the fit of the M1 model, especially for Spa-rea which had the lowest *B* (Figure 1D). However, both of these populations had a genome-wide significant influence of gBGC, and will still be considered in the following analyses. For all populations, M1* was not significantly better than M1, indicating either a lack of power for M1* or that the polarization error was negligible. The strength of gBGC (*B* = 4*N*_*e*_*b*) varied across GC centiles for all populations with Swe-sin and Kaz-sin showing the lowest standard error of the mean (0.009, Table 1, Figure 3A, B) and Ire-juv the highest (0.026). Because Ire-juv had the lowest number of SNPs per centile, it’s hard to disentangle sample-from biological variance but we note that Kaz-juv showed a similar standard error (0.025). The average value was overall congruent with what we observed in the analysis among populations (Table S2). We saw similar standard errors for the S→W mutation bias, λ (Table 1, Figure 3C, D).

**Table 1.** Population specific averages across GC centiles of λ, *B*, equilibrium GC content under mutational equilibrium alone, *GC*(1/[1+λ]), and when taking *B* into account *GC*(1/[1+λe^-*B*^]), and the observed GC content in the ancestral genome for the centile with the highest average pairwise difference *GC*(π_*max*_) and lowest density of coding sequence (*GC CDS*_*min*_). We also show standard error of the mean for λ and *B.*

| Population | $\lambda$ | $B$ | $GC$<br>$1/(1+\lambda)$ | $GC$<br>$1/(1+\lambda e^{-B})$ | $GC\ \pi_{max}$ | $GC$<br>$CDS_{min}$ |
| --- | --- | --- | --- | --- | --- | --- |
| Swe-sin | $2.99 \pm 0.010$ | $0.21 \pm 0.009$ | 0.25 | 0.29 | 0.27 | 0.35 |
| Spa-sin | $2.97 \pm 0.008$ | $0.21 \pm 0.010$ | 0.25 | 0.29 | 0.31 | 0.34 |
| Kaz-sin | $3.00 \pm 0.011$ | $0.20 \pm 0.009$ | 0.25 | 0.29 | 0.28 | 0.34 |
| Kaz-juv | $4.15 \pm 0.019$ | $0.79 \pm 0.025$ | 0.19 | 0.35 | 0.29 | 0.34 |
| Ire-juv | $3.54 \pm 0.027$ | $0.47 \pm 0.026$ | 0.22 | 0.31 | 0.29 | 0.34 |
| Spa-rea | $2.98 \pm 0.012$ | $0.16 \pm 0.011$ | 0.25 | 0.28 | 0.31 | 0.34 |

**Figure 3.**
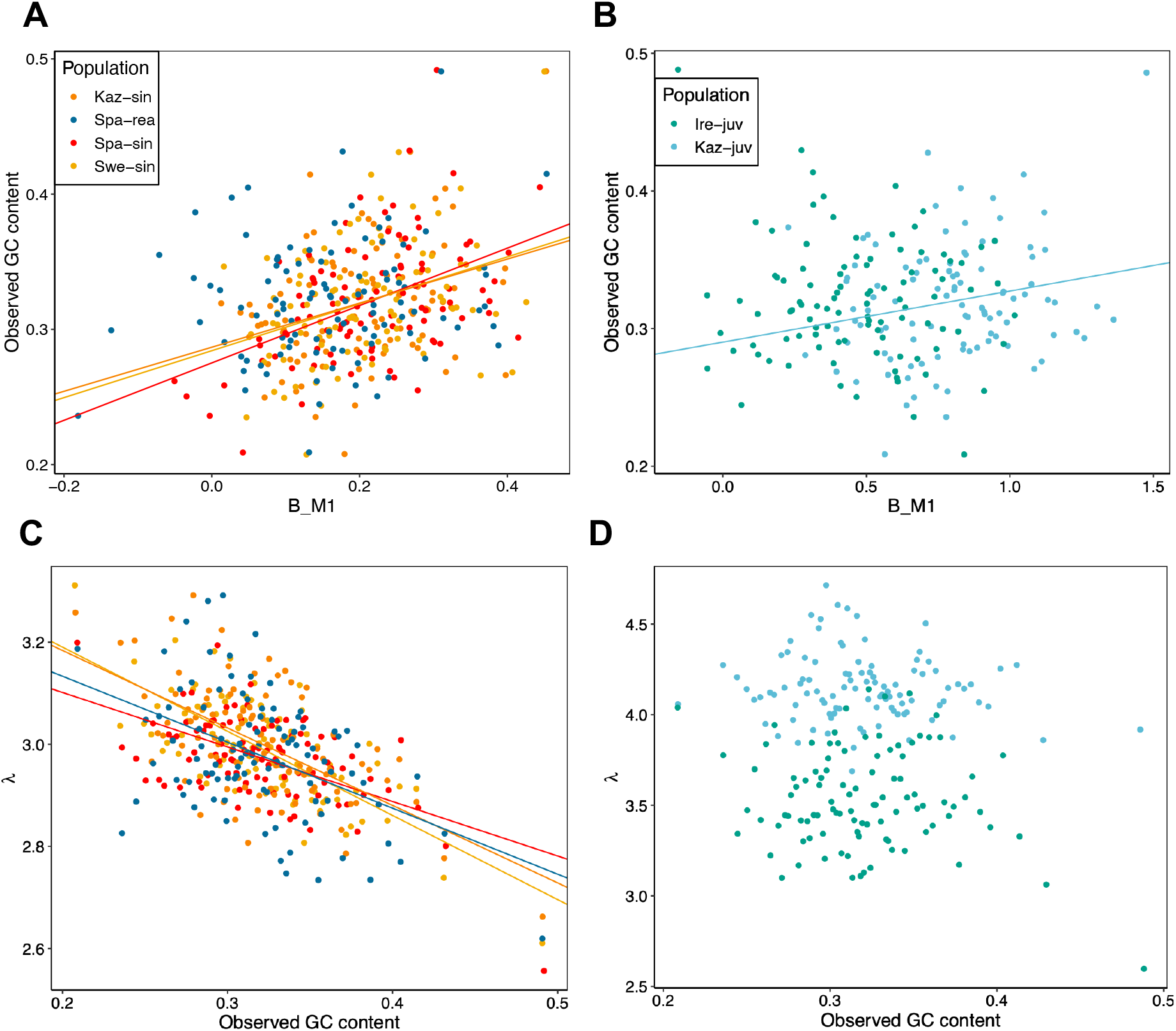
Relationship between *B,* λ and observed GC content in the ancestral genome. A) Association between *B* and observed GC content in the ancestral genome for the *L.-sinapis*-*L.reali* clade, and B) for the *L. juvernica* populations. Higher GC content was significantly consistent with greater *B* in all populations except Spa-rea and Ire-juv. C) Relationship between λ and GC content was negative for all populations in the *L.sinapis*-*L.reali* clade. D) Shows the same as C) but for the *L. juvernica* populations. Neither Kaz-juv nor Ire-juv showed significant associations between λ and GC content. Lines in plots represent significant linear regressions performed separately per population between the X- and Y variables.

To investigate the impact of variation in *N*_*e*_ across the genome we used genetic diversity, as a proxy for *N*_*e*_ and predictor of *B,* in separate linear regressions for each population (Figure S1A). Swe-sin and Kaz-sin showed significant negative relationships (*p* < 0.05), but limited variance explained (*R*^*2*^ ≈ 0.1 for both). The regressions were insignificant (*p* > 0.05) for the other populations (Figure S1A). Overall these results suggest that variation among centiles in *B* could be dominated by differences in conversion bias, *b*. An observation that supported this conclusion is that *B* significantly (*p* < 0.05; *R*^*2*^: 4-22 %) predicted GC content in four out of six populations (Figure 3A, B). Here GC content may serve as a proxy for recombination rate, assuming that differences in GC content has been caused by historically higher rates of recombination and thus stronger *B.* That two populations lacked a relationship with GC content may be explained partly by a lack of power for Ire-juv, which had the lowest number of SNPs per centile, while this explanation is less likely for Spa-rea. Nevertheless, for a majority of the populations considered here we saw a relationship between GC content in the ancestral genome and *B*, indicating that gBGC has been driving GC content evolution.

The mutation bias was significantly (*p* < 0.05, separate linear regression per population) negatively associated with observed GC content in the ancestral genome for all populations except Ire-juv and Kaz-juv (Figure 3C, D). To investigate if there was an association between λ and *B,* we performed separate linear regressions per population predicting λ with *B*. Higher estimates of λ across the genomes were consistent with larger values of *B* for all populations (*p* < 0.05) except Swe-sin and Spa-sin (Figure S1B). This indicates an inability of the model to separately estimate these parameters, or increased *B* in regions more prone to S→W mutations. The former explanation was unlikely given that the most common sign was negative in the regressions between λ and GC content.

### Mutation bias and gBGC influence the evolution of GC content

The equilibrium GC content in the presence of a S→W mutation bias, but in the absence of gBGC, can be calculated as 1/(1+λ) (Sueoka 1962). The observed GC content was higher than expected under mutational equilibrium alone across almost the entire genome for all populations (Figure 4A). When accounting for gBGC (1/(1+λe^-*B*^) (Li, et al. 1987; Bulmer 1991; Muyle, et al. 2011), the observed mean GC content was higher than the predicted equilibrium GC content in all populations except Kaz-juv (Table 1; Figure 4B).

**Figure 4.**
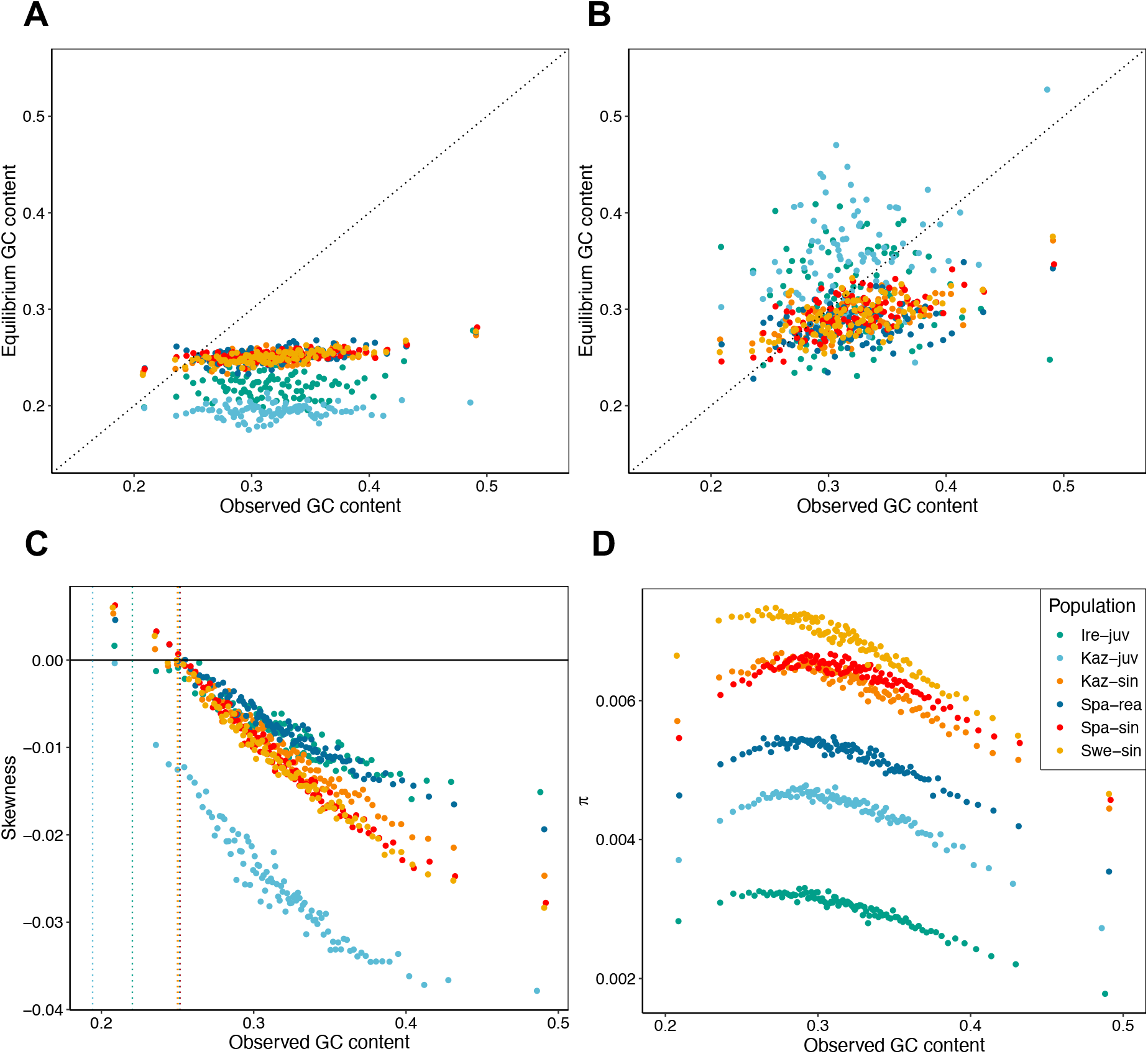
Observed GC content, equilibrium GC content and their association with λ, *B* and genetic diversity (π). A) Observed GC content compared to equilibrium GC content determined by mutation bias (λ) alone. B) Observed GC content compared to equilibrium GC content when accounting for gBGC. Dotted lines in (A) and (B) represent x = y. C) The skewness of the folded SFS shows the strong S→W bias in the segregating variation which increases with observed GC content in the ancestral genome. Extrapolating from the distribution of skewness values onto the y=0 line serves as a validation of the estimated λ. Dotted vertical lines represent the GC equilibrium under mutation bias alone, 1/(1+λ), for each population. D) The association between genetic diversity (*π*) and observed GC content. Points in all panels represent GC centiles.

Segregating variants hold information on the evolution of base composition. GC content will decrease if more S→W than W→S mutations reach fixation and vice versa. We can explore the fate of segregating variants by investigating the skewness of the folded site-frequency spectrum (SFS) (Figure 4C) (Glémin, et al. 2015). GC content is at equilibrium if skewness equals zero, evolves to higher GC content if the skew is positive and decreases if its negative. As expected from the relationship between observed and equilibrium GC content (Figure 4A), most of the centiles in all populations had a negative skew (Figure 4C).

### Pinnacle of genetic diversity close to GC equilibrium

We found a non-monotonic relationship between GC content and *π* (Figure 4D). The highest genetic diversity was observed close to the predicted genome-wide GC equilibrium, with diversity decreasing in both directions away from equilibrium GC content (Figure 4D). To test if this pattern could result from differential read coverage, we calculated the average read count per base pair in each GC centile per individual (Figure S2). Read coverage was generally even across most of the GC gradient except for a region around 35% GC where the *L. juvernica* populations show a signal consistent with a duplication event. Also, the centile with the greatest GC content showed high coverage in all populations. This is expected given the PCR bias against high and low GC regions in Illumina sequencing (e.g. Browne, et al. 2020). With the exception of *L. reali*, the GC content at the centile with the highest π, *GC*(π_max_), was at a level between the GC equilibrium defined by λ alone, *GC*(1/[1+λ]), and equilibrium when accounting for both λ and *B*, *GC*(1/[1+λe^-*B*^]). *GC*(π_max_) was lower for all populations than the GC content of the centile with the lowest density of coding sequence, *GC*(*CDS*_*min*_).

### The role of gBGC and mutation bias in shaping genetic diversity

Since gBGC mimics selection, the genetic diversity is directly dependent on the interaction between the strength of gBGC and the potential mutation bias (McVean and Charlesworth 1999; Glémin 2010). To understand how gBGC contributes to genetic diversity in *Leptidea,* we estimated the effects of gBGC and opposing mutation bias on genetic diversity by modelling the effect of *B* on the SFS (McVean and Charlesworth 1999). In the model, gBGC typically elevates the relative genetic diversity (π_*rel*_) compared to the case when gBGC is absent (*B* = 0) through increasing the equilibrium GC content. This allows for a greater influx of mutations as long as λ > 1 (Figure 5A). In *Leptidea*, genetic diversity (π) showed a non-monotonic relationship along the GC range (Figure 4D). In contrast, given values of λ around 3 and above, relevant for *Leptidea,* the model assuming gBGC-mutation-drift equilibrium (GMD) predicts a monotonic increase of π in the 0.2-0.5 GC range (Figure 5A). Using the output from the gBGC inference we could predict π_*rel*_ values for each GC centile and population from the GMD model (Figure 5B). The results showed that gBGC and mutation bias has the potential to elevate π compared to *B* = 0, by an average 2.6 % in Spa-rea, 3.3 % in Swe-sin and Kaz-sin, 3.5% in Spa-sin, 8 % in Ire-juv and 14.7 % in Kaz-juv.

**Figure 5:**
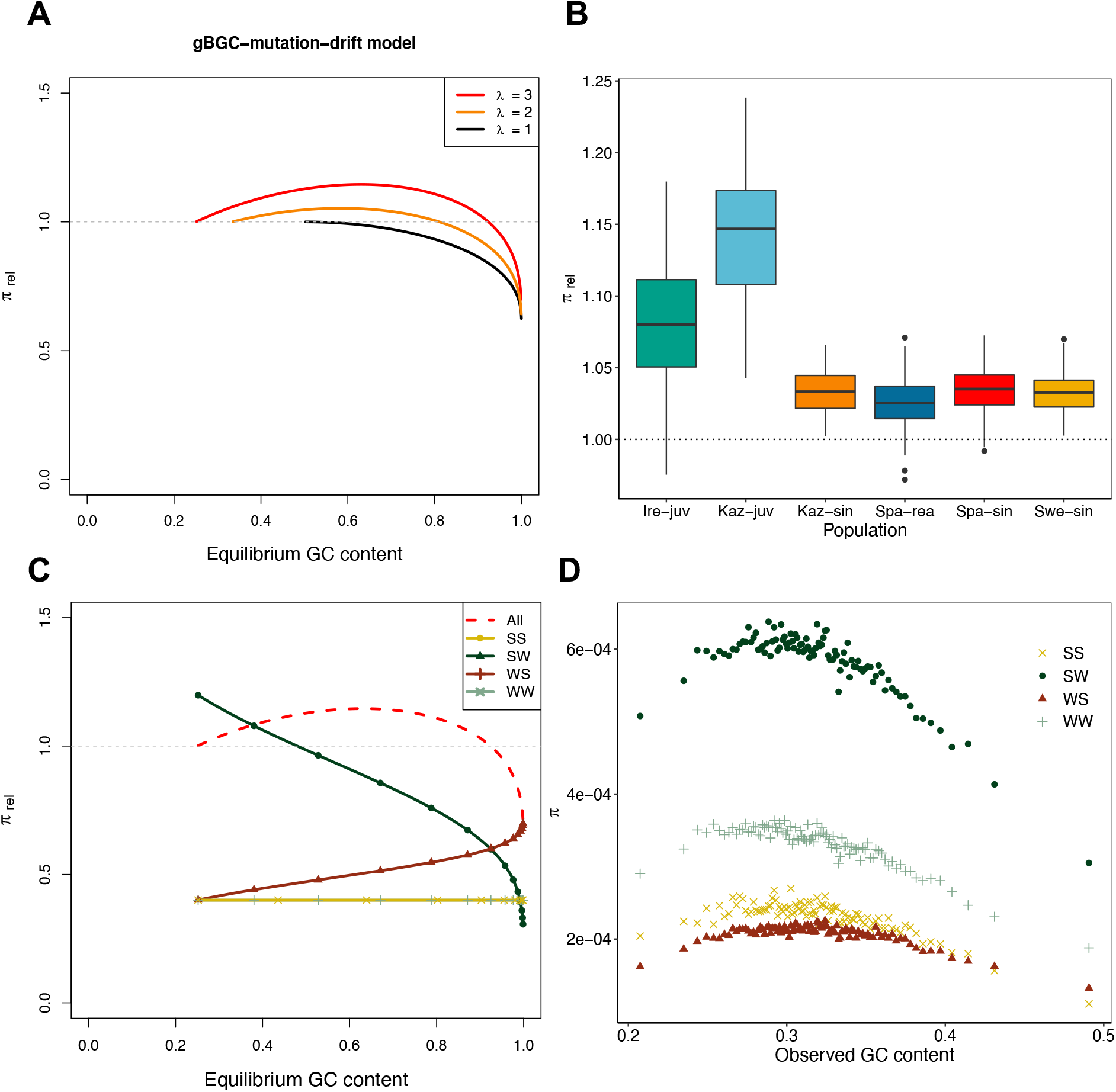
A model for genetic diversity under gBGC-mutation-drift equilibrium, predicted π_rel_ per population and π per mutation category. A) Genetic diversity relative to neutral (*B* = 0) across equilibrium GC content determined by *B* and λ. Lines begin at *B =* 0 and end at *B* = 8. The mutation bias is held constant. B) Genetic diversity values predicted from the gBGC-mutation-drift equilibrium model using output from the inference of gBGC. Most of the genome for each population have values of *B* and λ such that their relative strength boosts the long-term genetic diversity compared to *B* = 0. The lower and upper limit of the box correspond to the first and third quartiles. Upper and lower whiskers extend from the top- and bottom box limits to the largest/smallest value at maximum 1.5 times the inter-quartile range. C) Components of the gBGC mutation drift model. Only results from λ = 3 are shown. The separate mutation categories were standardized by mutational opportunity while “All” was standardized as in A). The genetic diversity is here assumed to be equal for N→N and W→S mutations (θ_N_ / θ_WS_ = 1). D) Genetic diversity in Swedish *L. sinapis* measured by average pairwise differences (π) across genomic GC content for all four mutation categories: S→S (SS), S→W (SW), W→S (WS), W→W (WW). The other populations are shown in Figure S3.

We can decompose the GMD model into four spectra standardized by their respective mutational opportunity (Figure 5C) to mimic the empirical data (Figure 5D). For example, the S→W category is standardized by equilibrium GC content. The four spectra include the GC-conservative/neutral spectra (W→W and S→S) and the GC-changing spectra (W→S and S→W) (Figure 5C). The contribution of GC-conservative mutation categories to π are unaffected by equilibrium GC content. In contrast, the influence of S→W on the SFS spectrum decreases as the intensity of *B* increases, and vice versa for W→S in the 0.2-0.5 GC range.

To understand the role gBGC plays in the variation of π with GC in *Leptidea*, we investigated the properties of the DAF spectra separately for all four mutation categories mentioned above. All mutation classes showed a qualitatively negative quadratic relationship between π and GC content (Figure 5D, Figure S3), which indicates that factors other than gBGC are the main determinants of the relationship between GC content and diversity (c.f. Figure 5C). A majority of the segregating sites were GC-changing and S→W contributed most to π across all centiles (Swe-sin: Figure 5D Others: Figure S3).

### The effects of linked selection and GC content on genetic diversity

Having rejected gBGC as a main contributor to the distribution of π along the GC gradient warrants the question: can the pattern be explained by reductions in diversity caused by linked selection? Linked selection has previously been shown to affect genetic diversity in butterfly genomes (Martin, et al. 2016; Talla, Soler, et al. 2019). Selection affecting linked sites will reduce genetic diversity unequally across the genome dependent on density of targets of selection and the rate of recombination. In agreement with this, CDS density, which can be used as a proxy for the intensity of linked selection in general but background selection in particular, was larger where π was lower (Figure 4D, Figure S4).

In addition, regional variation in mutation rate (μ) will also contribute to a heterogenous diversity landscape. We here suggest that GC content influences mutation rate for three reasons: i) π varies conspicuously with GC content (Figure 4D), ii) the S→W mutation bias appears to be affected by GC content (Figure 3 C), and, iii) GC content has been shown to be a major determinant of the mutation rate at CpG sites in humans (Fryxell and Moon 2005; Tyekucheva, et al. 2008; Schaibley, et al. 2013). Since guanine and cytosine are bound by three hydrogen bonds, one more than for adenine and thymine, it is believed that a higher local GC content reduces the formation of transient single-stranded states (Inman 1966). Cytosine deamination, which leads to C/G→T/A mutations, occurs at a higher rate in single-stranded DNA (Frederico, et al. 1993). Thus a higher GC content appears to reduce CpG mutation rates on a local scale of ca 2kb (Elango, et al. 2008). Mutation rate variation determined by local GC content outside the CpG context are less studied but negative correlations have been observed for most mutation classes in humans (Schaibley, et al. 2013).

To disentangle the relative contribution of GC content and CDS density on variation in π, we visualized the multivariate data by a coplot. The GC centiles were placed in five bins of equidistant GC content and separated by mutation category (Figure 6, Figure S4). The fifth bin was not considered as it included only a single centile with the highest GC content. First, we studied the association between GC content and CDS density (Figure 6A). GC content was negatively associated with CDS density in bin 1 and 2, while bin 3 showed no relationship and bin 4 a positive correlation (Figure 6A). Second, we considered the relationship between π and CDS density for all mutation categories. Here the general trend was negative, across GC bins, populations and mutation categories. In addition, the slopes got more negative with increasing GC content (Figure 6B, Figure S4).

**Figure 6:**
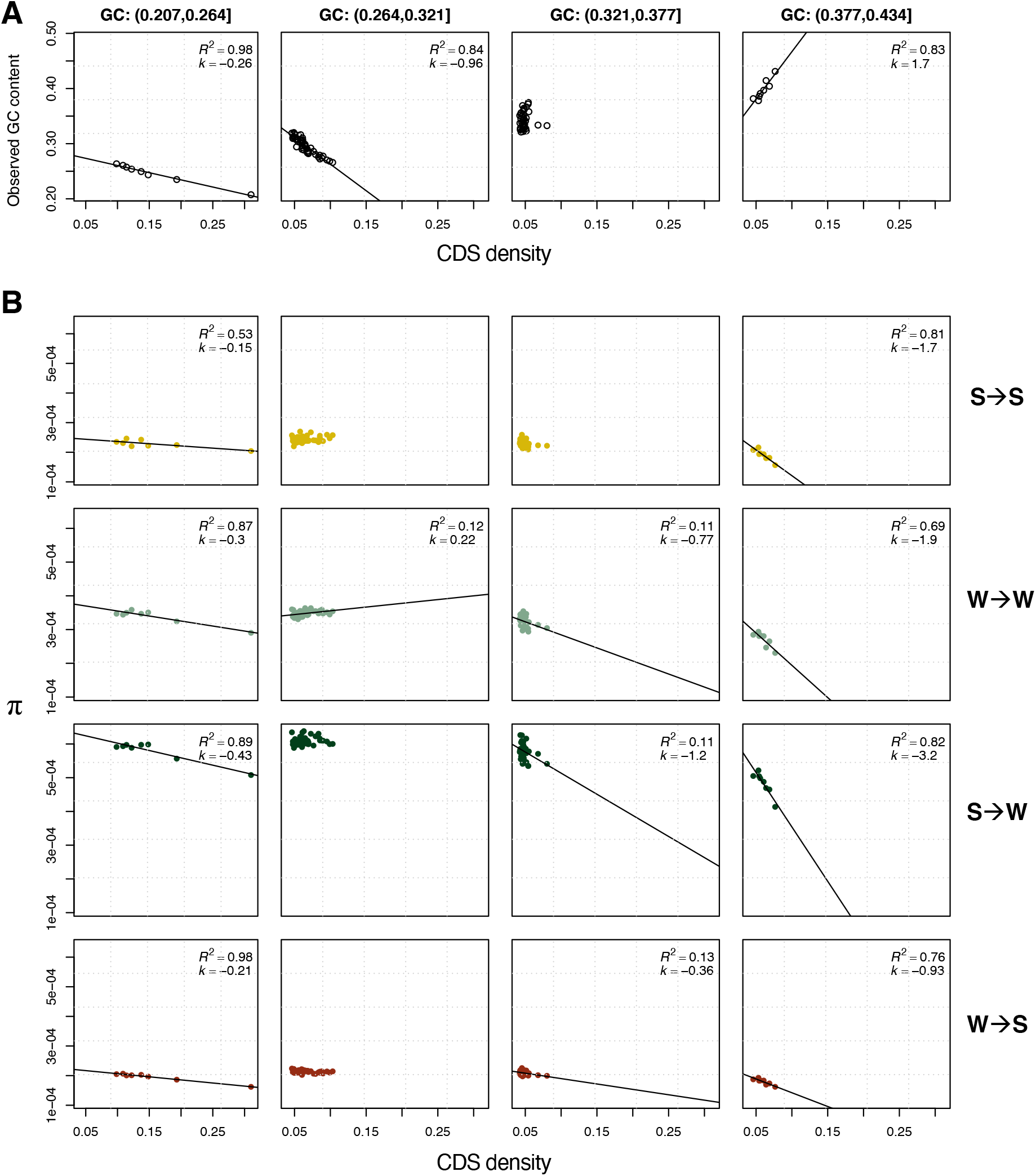
Relationship between π, CDS density and GC content. A) shows the relationship between CDS density and GC content for Swe-sin in four nonoverlapping equidistant intervals of GC content. B) instead shows the relationship between π and CDS density in the same bins separately for: S→S, W→W, S→W and W→S mutations. The fifth GC content bin is not shown because it includes only one centile. See Figure S4 for the other populations. *R*^*2*^ = proportion of variation explained, *k* = slope of regression (times 10^3^ for readability in B). GC bins 1-4 shown left to right. Mutation categories from top to bottom row: S→S, W→W, S→W and W→S.

For the GC-neutral mutation categories we observed the steepest negative slope when CDS density and GC content had a positive relationship (Bin 4, Figure 6, Figure S4). This may be caused by a joint effect of higher local GC content and CDS density contributing to a reduction in genetic diversity (Figure 6A, B). Despite a similar spread in CDS density, most populations showed fewer significant trends for bin 2. For Swe-sin the W→W mutation category even showed a positive slope (Figure 6B). Possibly a result of the negative relationship between GC content and CDS density giving an antagonistic response on diversity. When only GC content varied, π was also reduced for some but not all mutation categories and populations (Bin 3, Figure 6, Figure S4). When CDS density and GC content had a negative relationship, the slope was shallow but lower π was still consistent with a higher proportion of coding sequence (Bin 1, Figure 6). From these results we conclude that both GC content itself and linked selection affect diversity across the genome in *Leptidea* butterflies.

For the GC-changing mutation categories we observed patterns indicating that gBGC has affected genetic diversity either directly or indirectly (Figure 6B, Figure S4). The decomposed GMD model – with separate categories standardized for mutational opportunity – predicts that π will increase and decrease monotonically with GC content for W→S and S→W mutations, respectively (Figure 5C). Our results supported this conclusion with W→S mutations showing a shallower, and S→W a more pronounced negative slope compared to the GC-neutral mutation categories (Figure 6B, Figure S4). However, linked selection could interact with the distortion of the SFS caused by gBGC, which would constitute an indirect effect on π by gBGC. An argument against an indirect effect is that linked selection would be weaker or diminish were recombination is the highest, which most likely occur at greater GC content were *B* is stronger (see *Discussion*, Figure 3A, B) (Pouyet, et al. 2018). It is also possible that the W→S mutation rate is less restricted by high GC content as suggested by the negative relationship between λ and GC content for most populations (Figure 3C, D).

## Discussion

### The intensity of gBGC varies widely among species

In this study, we used whole-genome re-sequencing data from several populations of *Leptidea* butterflies to estimate gBGC and investigate its impact on rates and patterns of molecular evolution. Our data support previous observations that gBGC is present in butterflies (Galtier, et al. 2018). The genome-wide level of gBGC (*B*) varied from 0.17 - 0.80 among the investigated *Leptidea* populations. In general, *L. juvernica* populations had levels of *B* in line with previous estimates of gBGC in butterflies (0.69 - 1.16; Galtier, et al. 2018), while the other species had lower *B*, more in agreement with what has been observed in humans (0.38) (Glémin, et al. 2015).

### Determinants of gBGC variation in animals

Regression analysis suggested that the overall strength of gBGC among the *Leptidea* butterflies may depend more on interspecific variation in genome-wide recombination rate rather than differences in *N*_*e*_. Galtier et al. (2018) also showed a lack of correlation between *B* and longevity or propagule size (used as proxies for *N*_e_), across a wide sample of animals. We observed that chromosome number (a proxy for genome-wide recombination rate) was positively associated with *B* after excluding Spa-sin, which has recently experienced a change in karyotype. Galtier, et al. (2018) suggested that *B* may vary among species due to interspecific differences in transmission bias, *c*. This observation was supported by a study on honey bees (*Apis mellifera*) showing a substantial variation in transmission bias at non-crossover gene conversion events (0.10 – 0.15) among different subspecies (Kawakami, et al. 2019). Analyses of non-crossover gene conversion tracts in mice and humans showed that only conversion tracts including a single SNP were GC-biased (Li, et al. 2019). In contrast, the SNP closest to the end of a conversion tract determines the direction of conversion for all SNPs in a tract, in yeast (Lesecque, et al. 2013). Both these studies suggest that the impact of conversion tract length may be more complex than the multiplicative effect on conversion bias assumed in the *b* = *ncr* equation. The relative importance of recombination rate, transmission bias and conversion tract length, in divergence of *b* among populations and species remains to be elucidated.

### Butterfly population genomics in light of gBGC

Linkage maps for butterflies with high enough resolution to establish whether or not recombination is organized in hotspots is currently lacking (Davey, et al. 2016; Davey, et al. 2017; Halldorsson, et al. 2019). Nevertheless, recombination varies marginally (two-fold) between-but substantially within chromosomes in two species of the *Heliconius* genus (Davey, et al. 2017). Related to this, chromosome length is negatively correlated to both recombination rate and GC content in *H. melpomene* (Martin, et al. 2016; Davey, et al. 2017; Martin, et al. 2019), which is a pattern typical of gBGC (Pessia, et al. 2012). The higher GC content at fourfold degenerate (4D) sites on shorter chromosomes in *H. melpomene* was interpreted to be a consequence of stronger codon usage bias on short chromosomes (Martin, et al. 2016). An alternative explanation is that the higher recombination rate per base pair observed on smaller chromosomes leads to an increased intensity of gBGC and consequently a greater GC content. Galtier et al. (2018) showed significant positive correlation (*r* = 0.18-0.39) between GC content of the untranslated region and the third codon position in genes of three butterflies. This supports the conclusion that gBGC and possibly variation in mutation bias across the genome, affects codon usage evolution in butterflies. The degree of mutation bias in *H. melpomene* is unknown (as far as we know), but a λ ≈ 3 is possible given that *H. melpomene* has a genome-wide GC content of 32.8 % (Challis, et al. 2017), which is similar to the ancestral *Leptidea* genome and the *L. sinapis* reference assembly (Talla, et al. 2017; Talla, Johansson, et al. 2019). We conclude that assessment of natural selection using sequence data should also include disentangling the effects of potential confounding factors like gBGC, especially in taxa where this mechanism is prevalent (e.g. Bolívar, et al. 2016; Bolívar, et al. 2018; Pouyet, et al. 2018; Bolívar, et al. 2019).

### GC-biased gene conversion, mutation bias and genetic diversity

Many studies have in the recent decades investigated the association between genetic diversity and recombination rate and have in general found a positive relationship (e.g. Begun and Aquadro 1992; Nachman 1997; Kraft, et al. 1998; Cutter and Payseur 2003; Stevison and Noor 2010; Lohmueller, et al. 2011; Rao, et al. 2011; Langley, et al. 2012; Cutter and Payseur 2013; Mugal, et al. 2013; Corbett-Detig, et al. 2015; Wallberg, et al. 2015; Martin, et al. 2016; Pouyet, et al. 2018; Castellano, et al. 2019; Talla, Soler, et al. 2019). Somewhat later, debates on the determinants of so-called GC isochores in mammalian genomes gave rise to much research on the impact of gBGC on sequence evolution (Eyre-Walker 1999; Eyre-Walker and Hurst 2001; Meunier and Duret 2004; Duret, et al. 2006; reviewed in Duret and Galtier 2009). In this study we emphasize that gBGC and the widespread opposing mutation bias may also influence variation in genetic diversity across the genome. This can be considered as an extended neutral null model to which the importance of selective forces can be compared.

Several empirical studies have noted the impact of gBGC on genetic diversity. Castellano, et al. (2019) observed that the π of GC-changing mutations had a stronger positive correlation with recombination than GC-conservative mutations. Pouyet, et al. (2018) observed that in genomic regions with sufficiently high recombination to escape background selection, GC-neutral mutations were evolving neutrally while S→W mutations were disfavored and W→S mutations favored. This illustrates an important point that genomic regions where the diversity-reducing effects of background selection may be weakest or absent, are the same regions in which gBGC affects the SFS the most. Consequently, we suggest that future studies on the impact of linked selection also consider the impact of gBGC. A straight way forward would for example be to consider GC-neutral and GC-changing mutations separately (Castellano, et al. 2019).

The impact of gBGC on genetic diversity is dependent on the evolutionary timescale considered. For segregating variants, gBGC can only decrease diversity. If we also consider substitutions and model the evolution over longer timescales, gBGC may indirectly increase genetic diversity. In the GMD equilibrium model, gBGC raises genetic diversity indirectly by increasing GC content, which in turn allows greater mutational opportunity for S→W mutations. This can only be achieved when there is a S→W mutation bias greater than one and the intensity of gBGC is not too strong. Under identical conditions, gBGC may produce a positive correlation between recombination rate and genetic diversity through an increase in GC content. The impact of this effect will depend on the relative proportion of GC-neutral- and GC-changing variants. In the GMD model, the diversity of GC-neutral variants is unaffected by GC content. While this is a reasonable null model, it is also a simplistic view in light of the diversity-reducing effects on GC-neutral variants imposed by high GC content observed in our study. GC-neutral variants are only independent of gBGC on the timescale of segregating variation. Over longer timescales gBGC and mutation bias will cause GC-content to evolve towards an equilibrium which may or may not be conducive for GC-neutral mutations.

### Determinants of genetic diversity across the genome

Identifying determinants of genetic diversity and evaluating their relative importance remains a challenging task. First, we usually lack information on the relationship between GC content and mutation rate due to the sizable sequencing effort required to establish reliable estimates (Messer 2009). Divergence at synonymous sites have been used as a proxy for mutation rate (Martin, et al. 2016; Talla, Soler, et al. 2019), but synonymous divergence is a biased estimator of mutation rate in systems with *B* ≠ 0 (Bolívar, et al. 2016). In model organisms, such as humans, it has become feasible to study mutation rates using singletons in massive samples (>14,000 individuals; Schaibley, et al. 2013), or through large-scale sequencing of trios (Jónsson, et al. 2017). Second, the predictor variables of interest are often correlated (e.g. GC content and recombination rate in the presence of gBGC) which complicates interpretation for conventional multiple linear regression approaches (Talla, Soler, et al. 2019). A solution to this problem has been to use principal component regression (PCR) in which the PCs of predictor variables are used as regressors (Mugal, et al. 2013; Martin, et al. 2016; Dutoit, et al. 2017). Using this method, Dutoit, et al. (2017) found that the PC which explained most variation of π among 200 kb windows in the collared flycatcher genome was mainly composed of a negative correlation with GC. Martin, et al. (2016) considered 4D sites in *H. melpomene* and found that GC content was less important than gene density. It is likely that synonymous variants show greater impact of background selection compared to non-exonic variants, given the tight linkage between synonymous sites and nonsynonymous sites putatively under (purifying) selection. Instead of PCR we opted for an alternative approach in which the quadratic relationship between GC content and CDS density was binned into separate categories. Furthermore, by investigating the GC-neutral and GC-changing mutation categories separately, we could to some extent distinguish the effects of linked selection and GC content, from the effects of gBGC.

## Conclusion

In this study, we highlight that gBGC is a pervasive force, influencing rates and patterns of molecular evolution both among and across the genomes of *Leptidea* butterflies. We further emphasize that gBGC shapes genetic diversity and may – through fixation of W→S mutations – lead to a concomitant increase of diversity if opposed by a S→W mutation bias. This means that positive correlations between genetic diversity and recombination does not necessarily imply that selection is affecting diversity in the genome. Especially if the recombination rate is correlated with GC content, a pattern typical of gBGC. Here, we reject gBGC as a main determinant but recognizes its impact on diversity along with linked selection and GC content. Our model of how mutation bias and gBGC affects segregating variation provides a part of the puzzle linking the evolution of GC content to genetic diversity.

## Acknowledgements

This work was supported by a young investigator (VR 2013-4508) and a project research grant (VR 2019-4508) from the Swedish Research Council to NB. The authors acknowledge support from the National Genomics Infrastructure in Stockholm and Uppsala funded by the Science for Life Laboratory, the Knut and Alice Wallenberg Foundation and the Swedish Research Council, and SNIC/Uppsala Multidisciplinary Center for Advanced Computational Science for assistance with massively parallel sequencing, access to the UPPMAX computational infrastructure and the bioinformatics support team (WABI). The computations were performed on resources provided by the Swedish National Infrastructure for Computing (SNIC) at Uppsala University. We would also like to thank Per Unneberg, Venkat Talla, Karin Näsvall, Lars Höök, Daria Shipilina, Aleix Palahí Torres, Elenia Parkes, Yishu Zhu, Mahwash Jamy and Madeline Chase for helpful discussions regarding this work.

## Data accessibility

Raw sequence reads and binary alignment map files (.bam) have been deposited in the European Nucleotide Archive (ENA) under accession number PRJEB21838. In house developed scripts and pipelines are available at: xxx.

## Author contributions

NB and JB designed research. JB performed data analysis with input from NB and CFM. All authors approved the final version of the manuscript before submission.

## Notes

### Competing Interest Statement

The authors have declared no competing interest.

## References

Arbeithuber B, Betancourt AJ, Ebner T, Tiemann-Boege I. 2015. Crossovers are associated with mutation and biased gene conversion at recombination hotspots. Proceedings of the National Academy of Sciences of the United States of America 112:2109–2114.

Begun DJ, Aquadro CF. 1992. Levels of naturally occuring DNA polymorphism correlate with recombination rates in *D. melanogaster*. Nature 356:519–520.

Bewick AJ, Vogel KJ, Moore AJ, Schmitz RJ. 2017. Evolution of DNA methylation across insects. Molecular Biology and Evolution 34:654–665.

Birdsell JA. 2002. Integrating genomics, bioinformatics, and classical genetics to study the effects of recombination on genome evolution. Molecular Biology and Evolution 19:1181–1197.

Bolívar P, Guéguen L, Duret L, Ellegren H, Mugal CF. 2019. GC-biased gene conversion conceals the prediction of the nearly neutral theory in avian genomes. Genome Biology 20:5.

Bolívar P, Mugal CF, Nater A, Ellegren H. 2016. Recombination rate variation modulates gene sequence evolution mainly via GC-biased gene conversion, not Hill–Robertson interference, in an avian system. Molecular Biology and Evolution 33:216–227.

Bolívar P, Mugal CF, Rossi M, Nater A, Wang M, Dutoit L, Ellegren H. 2018. Biased inference of selection due to GC-biased gene conversion and the rate of protein evolution in flycatchers when accounting for it. Molecular Biology and Evolution 35:2475–2486.

Borges R, Szöllosi GJ, Kosiol C. 2019. Quantifying GC-biased gene conversion in great ape genomes using polymorphism-aware models. Genetics 212:1321–1336.

Brown TC, Jiricny J. 1987. A specific mismatch repair event protects mammalian cells from loss of 5-methylcytosine. Cell 50:945–950.

Browne PD, Nielsen TK, Kot W, Aggerholm A, Gilbert MTP, Puetz L, Rasmussen M, Zervas A, Hansen LH. 2020. GC bias affects genomic and metagenomic reconstructions, underrepresenting GC-poor organisms. GigaScience.

Bulmer M. 1991. The selection-mutation-drift theory of synonymous codon usage. Genetics 129:897–907.

Burri R, Nater A, Kawakami T, Mugal CF, Olason PI, Smeds L, Suh A, Dutoit L, Bureš S, Garamszegi LZ, et al. 2015. Linked selection and recombination rate variation drive the evolution of the genomic landscape of differentiation across the speciation continuum of Ficedula flycatchers. Genome Research 25:1656–1665.

Castellano D, Eyre-Walker A, Munch K. 2019. Impact of mutation rate and selection at linked sites on fine-scale DNA variation across the homininae genome. Genome Biology and Evolution 12:3550–3561.

Challis RJ, Kumar S, Dasmahapatra KK, Jiggins CD, Blaxter M. 2017. Lepbase - the lepidopteran genome database. BioRxiv [Online version].

Charlesworth B. 2012. The role of background selection in shaping patterns of molecular evolution and variation: evidence from variability on the *Drosophila* X chromosome. Genetics 191:233–246.

Charlesworth B, Morgan MT, Charlesworth D. 1993. The effect of deleterious mutations on neutral molecular variation. Genetics 134:1289–1303.

Chen JM, Cooper DN, Chuzhanova N, Férec C, Patrinos GP. 2007. Gene conversion: mechanisms, evolution and human disease. Nature Reviews Genetics 8:762–775.

Clément Y, Sarah G, Holtz Y, Homa F, Pointet S, Contreras S, Nabholz B, Sabot F, Sauné L, Ardisson M, et al. 2017. Evolutionary forces affecting synonymous variations in plant genomes. PLoS Genetics 13:e1006799–e1006799.

Comeron JM. 2017. Background selection as null hypothesis in population genomics: Insights and challenges from drosophila studies. Philosophical Transactions of the Royal Society B: Biological Sciences 372.

Coop G. 2016. Does linked selection explain the narrow range of genetic diversity across species? BioRxiv:042598-042598.

Corbett-Detig RB, Hartl DL, Sackton TB. 2015. Natural Selection Constrains Neutral Diversity across A Wide Range of Species. PLOS Biology 13:e1002112–e1002112.

Cutter AD, Payseur BA. 2013. Genomic signatures of selection at linked sites: unifying the disparity among species. Nature Reviews Genetics 14:262–274.

Cutter AD, Payseur BA. 2003. Selection at Linked Sites in the Partial Selfer Caenorhabditis elegans. Molecular Biology and Evolution 20:665–673.

Danecek P, Auton A, Abecasis G, Albers CA, Banks E, DePristo MA, Handsaker RE, Lunter G, Marth GT, Sherry ST, et al. 2011. The variant call format and VCFtools. Bioinformatics 27:2156–2158.

Davey JW, Barker SL, Rastas PM, Pinharanda A, Martin SH, Durbin R, McMillan WO, Merrill RM, Jiggins CD. 2017. No evidence for maintenance of a sympatric Heliconius species barrier by chromosomal inversions. Evolution Letters 1:138–154.

Davey JW, Chouteau M, Barker SL, Maroja L, Baxter SW, Simpson F, Joron M, Mallet J, Dasmahapatra KK, Jiggins CD. 2016. Major improvements to the *Heliconius melpomene* genome assembly used to confirm 10 chromosome fusion events in 6 Million years of butterfly evolution. G3: Genes, Genomes, Genetics 6:695–708.

Dincă V, Lukhtanov VA, Talavera G, Vila R. 2011. Unexpected layers of cryptic diversity in wood white *Leptidea* butterflies. Nature Communications 2:e324.

Dincă V, Wiklund C, Lukhtanov VA, Kodandaramaiah U, Noren K, Dapporto L, Wahlberg N, Vila R, Friberg M. 2013. Reproductive isolation and patterns of genetic differentiation in a cryptic butterfly species complex. Journal of Evolutionary Biology 26:2095–2106.

Duret L, Eyre-Walker A, Galtier N. 2006. A new perspective on isochore evolution. In: Elsevier. p. 71–74.

Duret L, Galtier N. 2009. Biased gene conversion and the evolution of mammalian genomic landscapes‥ Annual Review of Genomics and Human Genetics 10:285–311.

Duret L, Mouchiroud D. 1999. Expression pattern and, surprisingly, gene length shape codon usage in Caenorhabditis, Drosophila, and Arabidopsis. Proceedings of the National Academy of Sciences of the United States of America 96:4482–4487.

Dutoit L, Vijay N, Mugal CF, Bossu CM, Burri R, Wolf J, Ellegren H. 2017. Covariation in levels of nucleotide diversity in homologous regions of the avian genome long after completion of lineage sorting. Proceedings of the Royal Society B: Biological Sciences 284:20162756–20162756.

Elango N, Kim S-H, Vigoda E, Yi SV. 2008. Mutations of Different Molecular Origins Exhibit Contrasting Patterns of Regional Substitution Rate Variation. PLoS Computational Biology 4:e1000015–e1000015.

Eyre-Walker A. 1999. Evidence of selection on silent site base composition in mammals: Potential implications for the evolution of isochores and junk DNA. Genetics.

Eyre-Walker A, Hurst LD. 2001. The evolution of isochores. In: Nature Publishing Group. p. 549–555.

Eyre-Walker A, Keightley PD. 2007. The distribution of fitness effects of new mutations. In.

Eyre-Walker A, Woolfit M, Phelps T. 2006. The distribution of fitness effects of new deleterious amino acid mutations in humans. Genetics 173:891–900.

Felsenstein J. 1985. Phylogenies and the comparative method‥ The American Naturalist 125:1–15.

Frankham R. 1995. Effective population size/adult population size ratios in wildlife: A review. Genetical Research 66:95–107.

Frederico LA, Shaw BR, Kunkel TA. 1993. Cytosine Deamination in Mismatched Base Pairs. Biochemistry 32:6523–6530.

Fryxell KJ, Moon WJ. 2005. CpG mutation rates in the human genome are highly dependent on local GC content. Molecular Biology and Evolution 22:650–658.

Galtier N, Rousselle M. 2020. How Much Does Ne Vary Among Species? Genetics:genetics.303622.302020-genetics.303622.302020.

Galtier N, Roux C, Rousselle M, Romiguier J, Figuet E, Glemin S, Bierne N, Duret L. 2018. Codon usage bias in animals: disentangling the effects of natural selection, effective population size, and GC-biased gene conversion. Molecular Biology and Evolution 35:1092–1103.

Glémin S. 2010. Surprising fitness consequences of GC-biased gene conversion: I. mutation load and inbreeding depression. Genetics 185:939–959.

Glémin S, Arndt PF, Messer PW, Petrov D, Galtier N, Duret L. 2015. Quantification of GC-biased gene conversion in the human genome. Genome Research 25:1215–1228.

Halldorsson BV, Palsson G, Stefansson OA, Jonsson H, Hardarson MT, Eggertsson HP, Gunnarsson B, Oddsson A, Halldorsson GH, Zink F, et al. 2019. Human genetics: characterizing mutagenic effects of recombination through a sequence-level genetic map. Science 25:6425.

Hellmann I, Prüfer K, Ji H, Zody MC, Pääbo S, Ptak SE. 2005. Why do human diversity levels vary at a megabase scale? Genome Research 15:1222–1231.

Hodgkinson A, Eyre-Walker A. 2011. Variation in the mutation rate across mammalian genomes. In: Nature Publishing Group. p. 756–766.

Inman RB. 1966. A denaturation map of the λ phage DNA molecule determined by electron microscopy. Journal of Molecular Biology 18:464–476.

Jensen JD, Payseur BA, Stephan W, Aquadro CF, Lynch M, Charlesworth D, Charlesworth B. 2019. The importance of the neutral theory in 1968 and 50 years on: a response to Kern and Hahn 2018. Evolution 73:111–114.

Jones CM, Lim KS, Chapman JW, Bass C. 2018. Genome-wide characterization of DNA methylation in an invasive lepidopteran pest, the cotton bollworm *Helicoverpa armigera*. G3: Genes, Genomes, Genetics 8:779–787.

Jónsson H, Sulem P, Kehr B, Kristmundsdottir S, Zink F, Hjartarson E, Hardarson MT, Hjorleifsson KE, Eggertsson HP, Gudjonsson SA, et al. 2017. Parental influence on human germline de novo mutations in 1,548 trios from Iceland. Nature 549:519–522.

Kawakami T, Wallberg A, Olsson A, Wintermantel D, de Miranda JR, Allsopp M, Rundlöf M, Webster MT. 2019. Substantial Heritable Variation in Recombination Rate on Multiple Scales in Honeybees and Bumblebees. Genetics:genetics.302008.302019-genetics.302008.302019.

Kern AD, Hahn MW. 2018. The Neutral Theory in Light of Natural Selection. Molecular Biology and Evolution 35:1366–1371.

Kimura M. 1983. The Neutral Theory of Molecular Evolution. Cambridge: Cambridge University Press.

Kraft T, Säll T, Magnusson-Rading I, Nilsson N-O, Halldén C. 1998. Positive Correlation Between Recombination Rates and Levels of Genetic Variation in Natural Populations of Sea Beet (<em>Beta vulgaris</em> subsp. <em>maritima</em>). Genetics 150:1239.

Langley CH, Stevens K, Cardeno C, Lee YCG, Schrider DR, Pool JE, Langley SA, Suarez C, Corbett-Detig RB, Kolaczkowski B, et al. 2012. Genomic variation in natural populations of Drosophila melanogaster. Genetics.

Leal L, Talla V, Källman T, Friberg M, Wiklund C, Dincă V, Vila R, Backström N. 2018. Gene expression profiling across ontogenetic stages in the wood white (*Leptidea sinapis*) reveals pathways linked to butterfly diapause regulation. Molecular Ecology 27:935–948.

Lesecque Y, Mouchiroud D, Duret L. 2013. GC-Biased Gene Conversion in Yeast Is Specifically Associated with Crossovers: Molecular Mechanisms and Evolutionary Significance.

Lewontin RC. 1974. The genetic basis of evolutionary change. New York: Columbia Univ. Press.

Li R, Bitoun E, Altemose N, Davies RW, Davies B, Myers SR. 2019. A high-resolution map of non-crossover events reveals impacts of genetic diversity on mammalian meiotic recombination. Nature Communications 10:3900–3900.

Li WH, Tanimura M, Sharp PM. 1987. An evaluation of the molecular clock hypothesis using mammalian DNA sequences. Journal of Molecular Evolution 25:330–342.

Lohmueller KE, Albrechtsen A, Li Y, Kim SY, Korneliussen T, Vinckenbosch N, Tian G, Huerta-Sanchez E, Feder AF, Grarup N, et al. 2011. Natural Selection Affects Multiple Aspects of Genetic Variation at Putatively Neutral Sites across the Human Genome. PLoS Genetics 7:e1002326–e1002326.

Lukhtanov VA, Dincă V, Friberg M, Šíchová J, Olofsson M, Vila R, Marec F, Wiklund C. 2018. Versatility of multivalent orientation, inverted meiosis, and rescued fitness in holocentric chromosomal hybrids. Proceedings of the National Academy of Sciences of the United States of America 115:E9610–E9619.

Lukhtanov VA, Dincă V, Friberg M, Vila R, Wiklund C. 2020. Incomplete Sterility of Chromosomal Hybrids: Implications for Karyotype Evolution and Homoploid Hybrid Speciation. Frontiers in Genetics 11:1205–1205.

Lukhtanov VA, Dincă V, Talavera G, Vila R. 2011. Unprecedented within-species chromosome number cline in the wood white butterfly *Leptidea sinapis* and its significance for karyotype evolution and speciation. BMC Evolutionary Biology 11:e109.

Lynch M. 2007. The origins of genome architecture. Sunderland, MA: Sinauer Associates.

Lynch M, Ackerman MS, Gout JF, Long H, Sung W, Thomas WK, Foster PL. 2016. Genetic drift, selection and the evolution of the mutation rate. In: Nature Publishing Group. p. 704–714.

Mackintosh A, Laetsch DR, Hayward A, Charlesworth B, Waterfall M, Vila R, Lohse K. 2019. The determinants of genetic diversity in butterflies. Nature Communications 10:3466.

Maeda T. 1939. Chiasma studies in the silkworm, *Bombyx mori* L. Japanese Journal of Genetics 15:118–127.

Mancera E, Bourgon R, Brozzi A, Huber W, Steinmetz LM. 2008. High-resolution mapping of meiotic crossovers and non-crossovers in yeast. Nature 454:479–485.

Marais G. (pdf00327 co-authors). 2003. Biased gene conversion: implications for genome and sex evolution. Trends in Genetics 19:330–338.

Martin SH, Davey JW, Salazar C, Jiggins CD. 2019. Recombination rate variation shapes barriers to introgression across butterfly genomes. PLOS Biology 17:e2006288.

Martin SH, Möst M, Palmer WJ, Salazar C, McMillan WO, Jiggins FM, Jiggins CD. 2016. Natural selection and genetic diversity in the butterfly *Heliconius melpomene*. Genetics 203:525–541.

Maynard Smith J, Haigh J. 1974. The hitch-hiking effect of a favourable gene. Genetical Research 23:23–35.

McVean GAT, Charlesworth B. 1999. A population genetic model for the evolution of synonymous codon usage: patterns and predictions. Genetical Research 74:145–158.

Messer PW. 2009. Measuring the rates of spontaneous mutation from deep and large-scale polymorphism data. Genetics 182:1219–1232.

Meunier J, Duret L. 2004. Recombination drives the evolution of GC-content in the human genome. Molecular Biology and Evolution.

Mugal CF, Nabholz B, Ellegren H. 2013. Genome-wide analysis in chicken reveals that local levels of genetic diversity are mainly governed by the rate of recombination. BMC Genomics 14:e86.

Mugal CF, Weber CC, Ellegren H. 2015. GC-biased gene conversion links the recombination landscape and demography to genomic base composition: GC-biased gene conversion drives genomic base composition across a wide range of species. Bioessays 37:1317–1326.

Muyle A, Serres-Giardi L, Ressayre A, Escobar J, Glémin S. 2011. GC-biased gene conversion and selection affect GC content in the *Oryza* genus (rice). Molecular Biology and Evolution 28:2695–2706.

Nachman MW. 1997. Patterns of DNA Variability at X-Linked Loci in Mus domesticus. Genetics 147.

Nagylaki T. 1983a. Evolution of a finite population under gene conversion. Proceedings of the National Academy of Sciences of the United States of America 80:6278–6281.

Nagylaki T. 1983b. Evolution of a large population under gene conversion. Proceedings of the National Academy of Sciences of the United States of America 80:5941–5945.

Nevo E, Beiles A, Ben-Shlomo R editors. Evolutionary Dynamics of Genetic Diversity. 1984 1984//: Berlin, Heidelberg.

Paradis E, Schliep K. 2018. Ape 5.0: an environment for modern phylogenetics and evolutionary analyses in R. Bioinformatics 35:526–528.

Perry J, Ashworth A. 1999. Evolutionary rate of a gene affected by chromosomal position. Current Biology 9:987–989.

Pessia E, Popa A, Mousset S, Rezvoy C, Duret L, Marais GA. 2012. Evidence for widespread GC-biased gene conversion in eukaryotes. Genome Biology and Evolution 4:675–682.

Pouyet F, Aeschbacher S, Thiéry A, Excoffier L. 2018. Background selection and biased gene conversion affect more than 95% of the human genome and bias demographic inferences. eLife 7:e36317.

Provataris P, Meusemann K, Niehuis O, Grath S, Misof B. 2018. Signatures of DNA methylation across insects suggest reduced DNA methylation levels in Holometabola. Genome Biology and Evolution 10:1185–1197.

Quinlan AR, Hall IM. 2010. BEDTools: a flexible suite of utilities for comparing genomic features. Bioinformatics 26:841–842.

R Core Team T. 2020. R: A language and environment for statistical computing. Vienna, Austria: Foundation for Statistical Computing.

Rao Y, Sun L, Nie Q, Zhang X. 2011. The influence of recombination on SNP diversity in chickens. Hereditas.

Rettelbach A, Nater A, Ellegren H. 2019. How linked selection shapes the diversity landscape in *Ficedula* flycatchers. Genetics 212:277–285.

Romiguier J, Gayral P, Ballenghien M, Bernard A, Cahais V, Chenuil A, Chiari Y, Dernat R, Duret L, Faivre N, et al. 2014. Comparative population genomics in animals uncovers the determinants of genetic diversity. Nature 515:261–263.

Roy AM, Carroll ML, Nguyen SV, Salem AH, Oldridge M, Wilkie AOM, Batzer MA, Deininger PL. 2000. Potential gene conversion and source genes for recently integrated Alu elements. Genome Research 10:1485–1495.

Schaibley VM, Zawistowski M, Wegmann D, Ehm MG, Nelson MR, St.Jean PL, Abecasis GR, Novembre J, Zöllner S, Li JZ. 2013. The influence of genomic context on mutation patterns in the human genome inferred from rare variants. Genome Research 23:1974–1984.

Sexton CE, Han MV. 2019. Paired-end mappability of transposable elements in the human genome. Mobile DNA 10:29–29.

Šíchová J, Voleníková A, Dincə V, Nguyen P, Vila R, Sahara K, Marec F. 2015. Dynamic karyotype evolution and unique sex determination systems in Leptidea wood white butterflies Speciation and evolutionary genetics. BMC Evolutionary Biology 15.

Smeds L, Mugal CF, Qvarnström A, Ellegren H. 2016. High-Resolution Mapping of Crossover and Non-crossover Recombination Events by Whole-Genome Re-sequencing of an Avian Pedigree. PLoS Genetics 12:e1006044–e1006044.

Smith TCA, Arndt PF, Eyre-Walker A. 2018. Large scale variation in the rate of germ-line de novo mutation, base composition, divergence and diversity in humans. PLoS Genetics 14:e1007254–e1007254.

Spencer CC, Deloukas P, Hunt S, Mullikin J, Myers S, Silverman B, Donnelly P, Bentley D, McVean G. 2006. The influence of recombination on human genetic diversity. PLoS Genetics 2:e148.

Stevison LS, Noor MAF. 2010. Genetic and Evolutionary Correlates of Fine-Scale Recombination Rate Variation in Drosophila persimilis. Journal of Molecular Evolution 71:332–345.

Sueoka N. 1962. On the genetic basis of variation and heterogeneity of DNA base composition. Proceedings of the National Academy of Sciences of the United States of America 48:582–592.

Suomalainen E, Cook LM, Turner JRG. 1973. Achiasmatic oogenesis in the *Heliconiine* butterflies. Hereditas 74:302–304.

Talla V, Johansson A, Dincă V, Vila R, Friberg M, Wiklund C, Backström N. 2019. Lack of gene flow: narrow and dispersed differentiation islands in a triplet of *Leptidea* butterfly species. Molecular Ecology 28:3756–3770.

Talla V, Soler L, Kawakami T, Dincă V, Vila R, Friberg M, Wiklund C, Backström N. 2019. Dissecting the effects of selection and mutation on genetic diversity in three wood white (*Leptidea* sp.) species. Genome Biology and Evolution 11:2875–2886.

Talla V, Suh A, Kalsoom F, Dincă V, Vila R, Friberg M, Wiklund C, Backström N. 2017. Rapid increase in genome size as a consequence of transposable element hyperactivity in wood-white (*Leptidea*) butterflies. Genome Biology and Evolution 9:2491–2505.

Torres R, Stetter MG, Hernandez RD, Ross-Ibarra J. 2020. The Temporal Dynamics of Background Selection in Non-equilibrium Populations. Genetics:genetics.302892.302019-genetics.302892.302019.

Turner JRG, Sheppard PM. 1975. Absence of crossing-over in female butterflies (*Heliconius*). Heredity 34:265–269.

Tyekucheva S, Makova KD, Karro JE, Hardison RC, Miller W, Chiaromonte F. 2008. Human-macaque comparisons illuminate variation in neutral substitution rates. Genome Biology 9:R76–R76.

Wallberg A, Glémin S, Webster MT. 2015. Extreme Recombination Frequencies Shape Genome Variation and Evolution in the Honeybee, Apis mellifera. PLoS Genetics 11:e1005189–e1005189.

Wickham H. 2016. Ggplot2: Elegant Graphics for Data Analysis. New York: Springer Verlag. Wolfram Research I. 2019. Mathematica. Champaign, Illinois.

