## Supplementary Information for "The effects of GC-biased gene conversion on patterns of genetic diversity among and across butterfly genomes"

**Table S1:** Individuals used to polarize the SNPs and number of polarized SNPs for each polarization, filter set, analysis and population. The abbreviations for the filter sets are: NE = all non-exonic sites separated into "strict" ($NE_S$) and "liberal" ($NE_L$) polarization schemes, NE CpG- = no ancestral CpG-prone sites, NE CpG+ = ancestral CpG-prone sites. Polarization abbreviations are: s = *L. sinapis*, j = *L. juvernica*, r = *L. reali*, kj = Kazakhstani *L. juvernica* individuals, ij = Irish *L. juvernica* individuals, HD = highest mean read-depth.

| | ***L. juvernica*** | | ***L. reali*** | ***L. sinapis*** | | |
|---|---|---|---|---|---|---|
| **Population** | Ire-juv | Kaz-juv | Spa-rea | Swe-sin | Spa-sin | Kaz-sin |
| **Strict polarization** | 30s10r10kj | 30s10r10ij | 30s20j | 10r20j | 10r20j | 10r20j |
| **$NE_S$** | 266,103 | 766,700 | 1,252,453 | 1,993,236 | 2,114,098 | 1,700,701 |
| **NE CpG-** | 177,527 | 511,772 | 829,753 | 1,307,791 | 1,388,478 | 1,115,788 |
| **NE CpG+** | 88,576 | 254,928 | 422,700 | 685,445 | 725,620 | 584,913 |
| **GC centile** | 2,661 | 7,667 | 12,524 | 19,932 | 21,140 | 17,007 |
| **Liberal polarization** | 1s1r HD | 1s1r HD | 1s1j HD | 1r1j HD | 1r1j HD | 1r1j HD |
| **$NE_L$** | 2,108,580 | 3,251,996 | 2,663,362 | 3,826,751 | 3,765,982 | 3,339,597 |

**Table S2.**

Statistics from the gBGC maximum likelihood estimation. Estimates of *B* and λ from the M1 and M1* models and polarization error proportion for W→S (eWS) and S→W (eSW) polarized SNP. Also shown is the p-value for the likelihood ratio tests between the M1 and M1* model. Values are rounded to four digits. The abbreviations for the filter sets are: NE = all non-exonic sites separated into "strict" ($NE_S$) and "liberal" ($NE_L$) polarization schemes, NE CpG- = all non-genic, non-ancestral CpG sites, NE CpG+ = all non-genic ancestral CpG sites.

| Population | Filter | *B* M1 | *B* M1* | λ M1 | λ M1* | eWS | eSW | *p*-value (LRT) |
|---|---|---|---|---|---|---|---|---|
| Swe-sin | $NE_S$ | 0.2103 | 0.2184 | 2.9603 | 2.9614 | 0.0007 | 0.0000 | 0.4120 |
| Spa-sin | $NE_S$ | 0.2214 | 0.2417 | 2.9531 | 2.9362 | 0.0202 | 0.0121 | 0.0002 |
| Kaz-sin | $NE_S$ | 0.2083 | 0.2265 | 2.9703 | 2.9476 | 0.0241 | 0.0142 | 0.0215 |
| Kaz-juv | $NE_S$ | 0.7792 | 0.7932 | 4.0847 | 4.0899 | 0.0003 | 0.0000 | 0.0287 |
| Ire-juv | $NE_S$ | 0.4597 | 0.5403 | 3.4839 | 3.5165 | 0.0055 | 0.0000 | 0.0000 |
| Spa-rea | $NE_S$ | 0.1661 | 0.2109 | 2.9570 | 2.9611 | 0.0052 | 0.0000 | 0.0000 |
| Swe-sin | $NE_S$ CpG- | 0.1855 | 0.1890 | 2.6815 | 2.7121 | 0.0084 | 0.0192 | 0.9579 |
| Spa-sin | $NE_S$ CpG- | 0.1970 | 0.2151 | 2.6844 | 2.7262 | 0.0104 | 0.0233 | 0.0000 |
| Kaz-sin | $NE_S$ CpG- | 0.1690 | 0.1777 | 2.6752 | 2.7204 | 0.0125 | 0.0277 | 0.5405 |
| Kaz-juv | $NE_S$ CpG- | 1.2010 | 1.2240 | 4.2590 | 4.2687 | 0.0003 | 0.0000 | 0.0032 |
| Ire-juv | $NE_S$ CpG- | 0.6771 | 0.7902 | 3.4324 | 3.5023 | 0.0086 | 0.0112 | 0.0000 |
| Spa-rea | $NE_S$ CpG- | 0.1721 | 0.2224 | 2.7128 | 2.7244 | 0.0032 | 0.0000 | 0.0004 |
| Swe-sin | $NE_S$ CpG+ | 0.2122 | 0.2212 | 3.3593 | 3.3523 | 0.0035 | 0.0001 | 0.4478 |
| Spa-sin | $NE_S$ CpG+ | 0.2239 | 0.2331 | 3.3490 | 3.3384 | 0.0050 | 0.0002 | 0.2230 |
| Kaz-sin | $NE_S$ CpG+ | 0.2382 | 0.2629 | 3.4054 | 3.3898 | 0.0084 | 0.0001 | 0.0462 |
| Kaz-juv | $NE_S$ CpG+ | 0.4864 | 0.4931 | 4.1653 | 4.1591 | 0.0024 | 0.0001 | 0.7180 |
| Ire-juv | $NE_S$ CpG+ | 0.3617 | 0.4340 | 3.9744 | 3.9548 | 0.0180 | 0.0000 | 0.0830 |
| Spa-rea | $NE_S$ CpG+ | 0.1528 | 0.1900 | 3.3866 | 3.3607 | 0.0137 | 0.0000 | 0.1250 |
| Swe-sin | $NE_L$ | 0.1984 | 0.2071 | 2.9298 | 2.9288 | 0.0015 | 0.0000 | 0.2471 |
| Spa-sin | $NE_L$ | 0.2499 | 0.2767 | 2.9668 | 2.9214 | 0.0489 | 0.0308 | 0.0000 |
| Kaz-sin | $NE_L$ | 0.1976 | 0.2190 | 2.9560 | 2.8672 | 0.0749 | 0.0467 | 0.0017 |
| Kaz-juv | $NE_L$ | 0.4828 | 0.6878 | 3.4269 | 3.4006 | 0.0350 | 0.0000 | 0.0000 |
| Ire-juv | $NE_L$ | 0.1795 | 0.3427 | 2.9393 | 2.9091 | 0.0345 | 0.0000 | 0.0000 |
| Spa-rea | $NE_L$ | 0.2154 | 0.2729 | 2.9553 | 2.9501 | 0.0099 | 0.0000 | 0.0000 |

**Figure S1.**

Linear regressions between parameters from the investigation of *B* and λ in GC centiles. Points represents GC centiles. A) Relationship between *B* and π. B) Relationship between the estimated λ and *B*. Only significant ($p < 0.05$) regression lines are shown.

**A** **B**

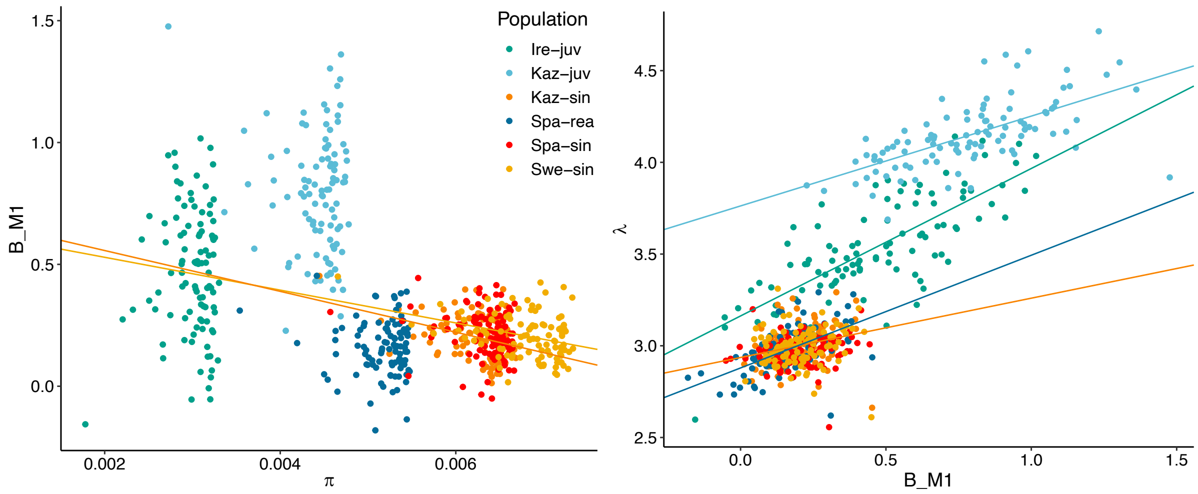

**Figure S2.**

Coverage per centile, individual and population. Fitted lines are curves from generalized additive models with integrated smoothness (settings for geom_smooth (Wickham 2016): method = 'gam' and formula 'y ~ s(x, bs = "cs")).

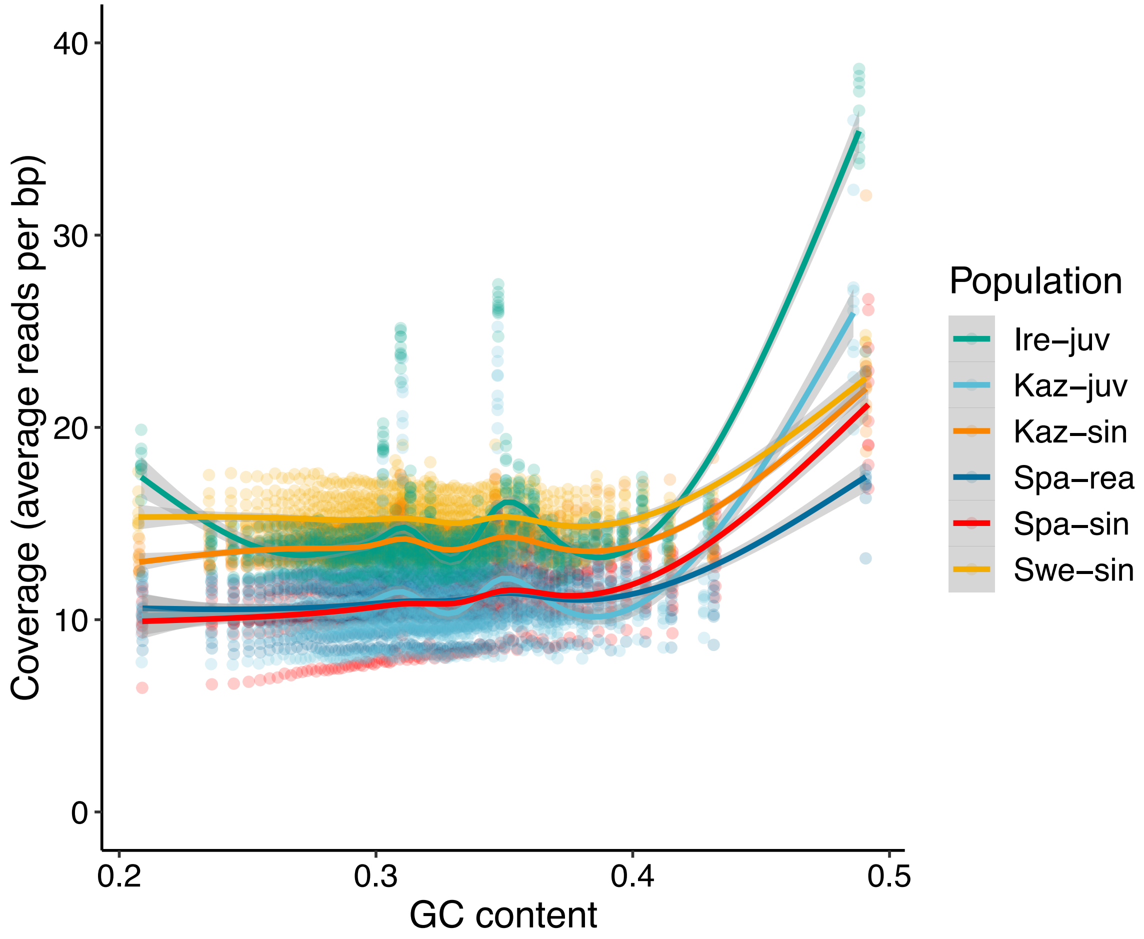

**Figure S3.**

Genetic diversity, π, across genomic GC content per mutation class: S→S (SS), S→W (SW), W→S (WS) and W→W (WW), for all populations except Swe-sin (see *Main text*).

**Spa-sin**

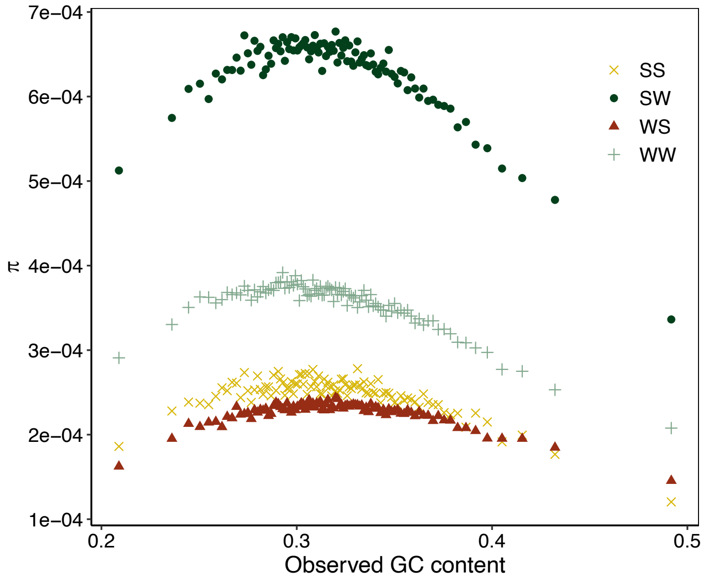


**Kaz-sin**

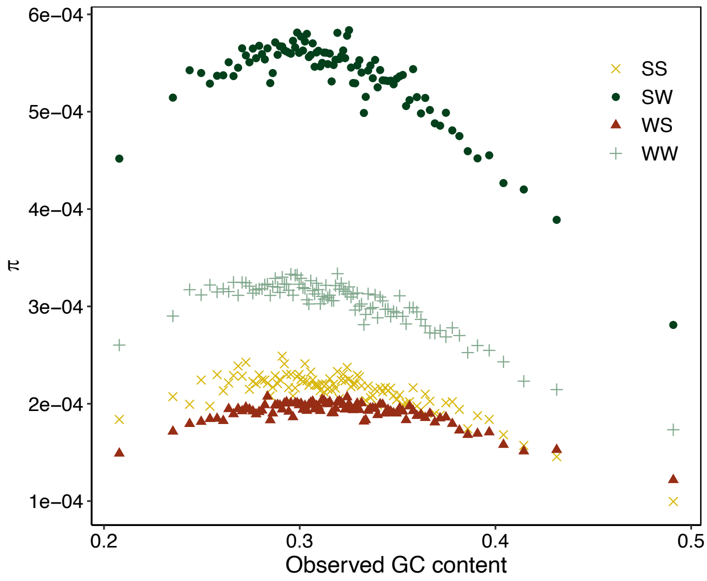


**Kaz-juv**

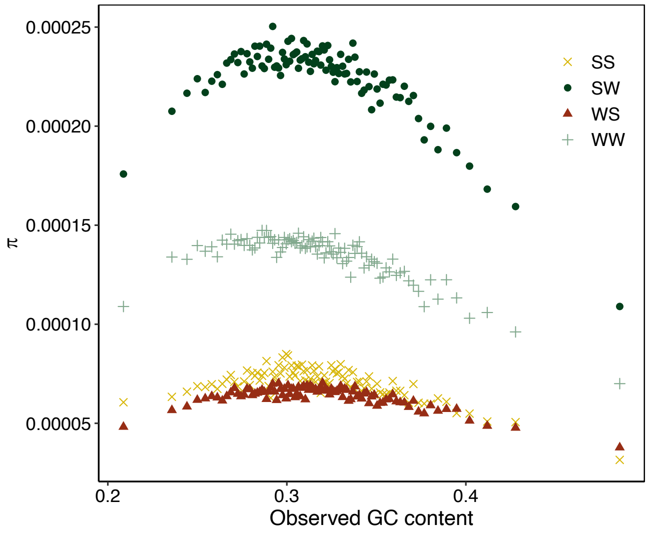


**Ire-juv**

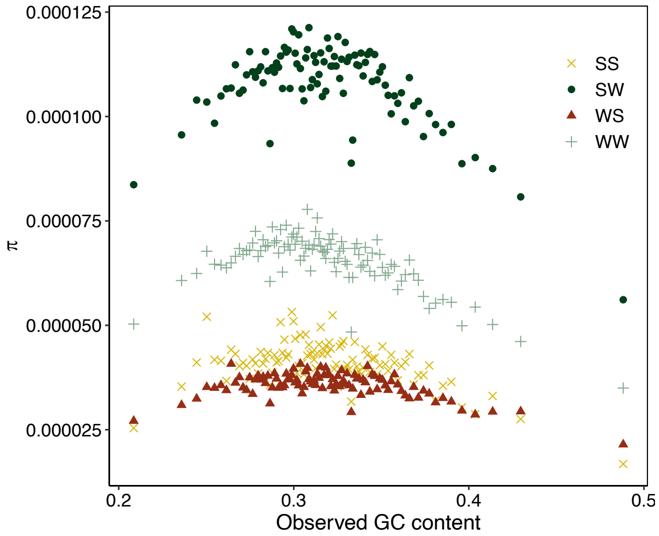


**Spa-rea**

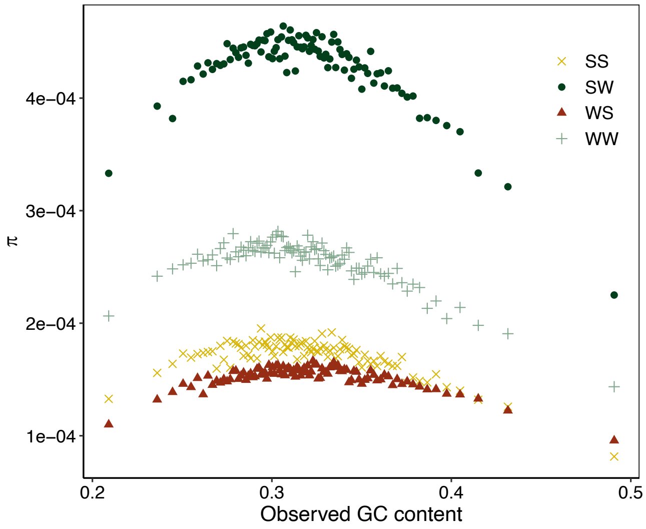

**Figure S4.**

Relationship between π, CDS density and GC content for all populations except Swe-sin (see *Main text*). $R^2$ = proportion of variation explained, $k$ = slope of regression (times $10^3$ for readability in B). GC bins 1-4 shown left to right. Mutation categories from top to bottom row: S→S, W→W, S→W and W→S.

**Spa-sin**

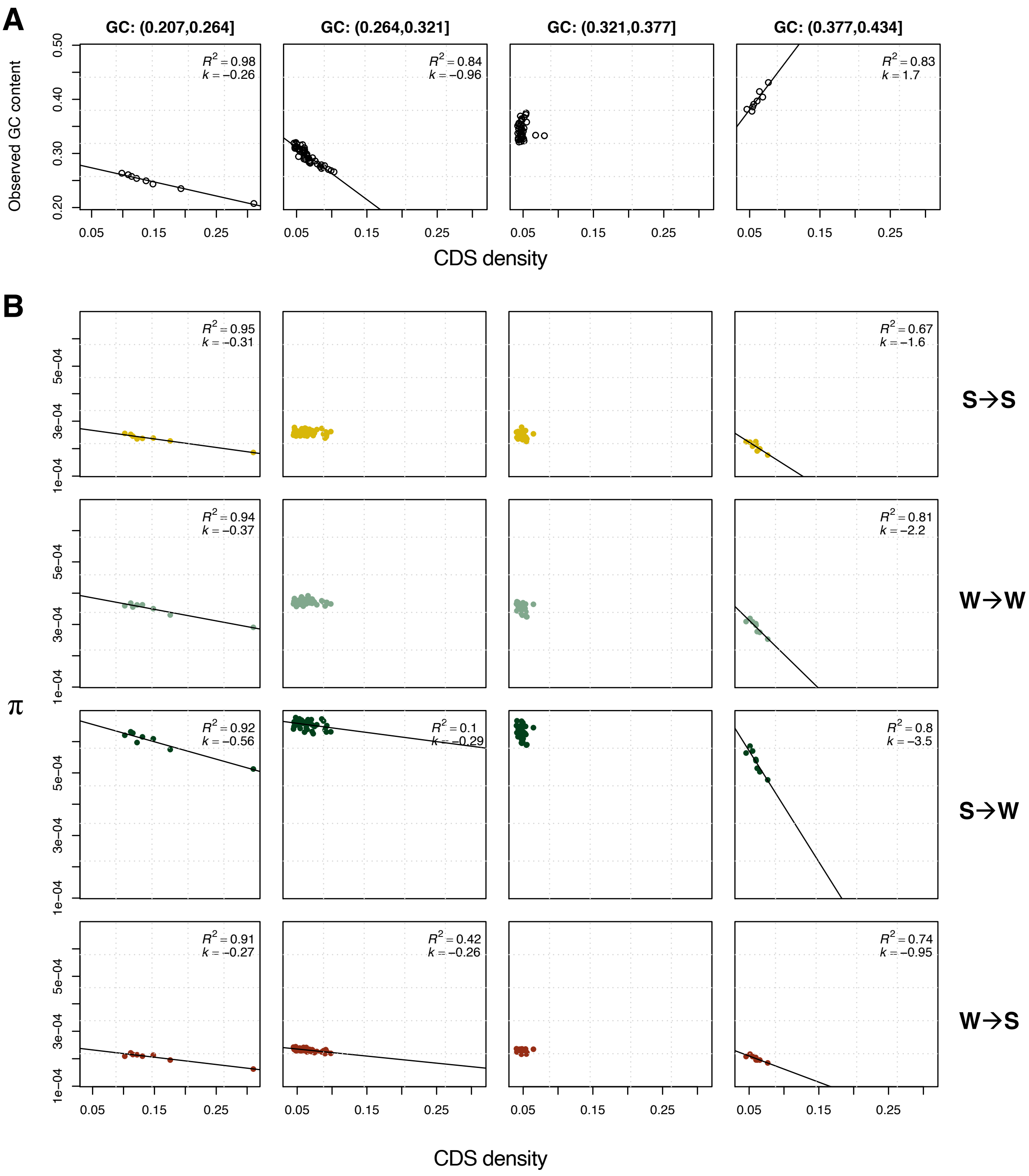

**Kaz-sin**

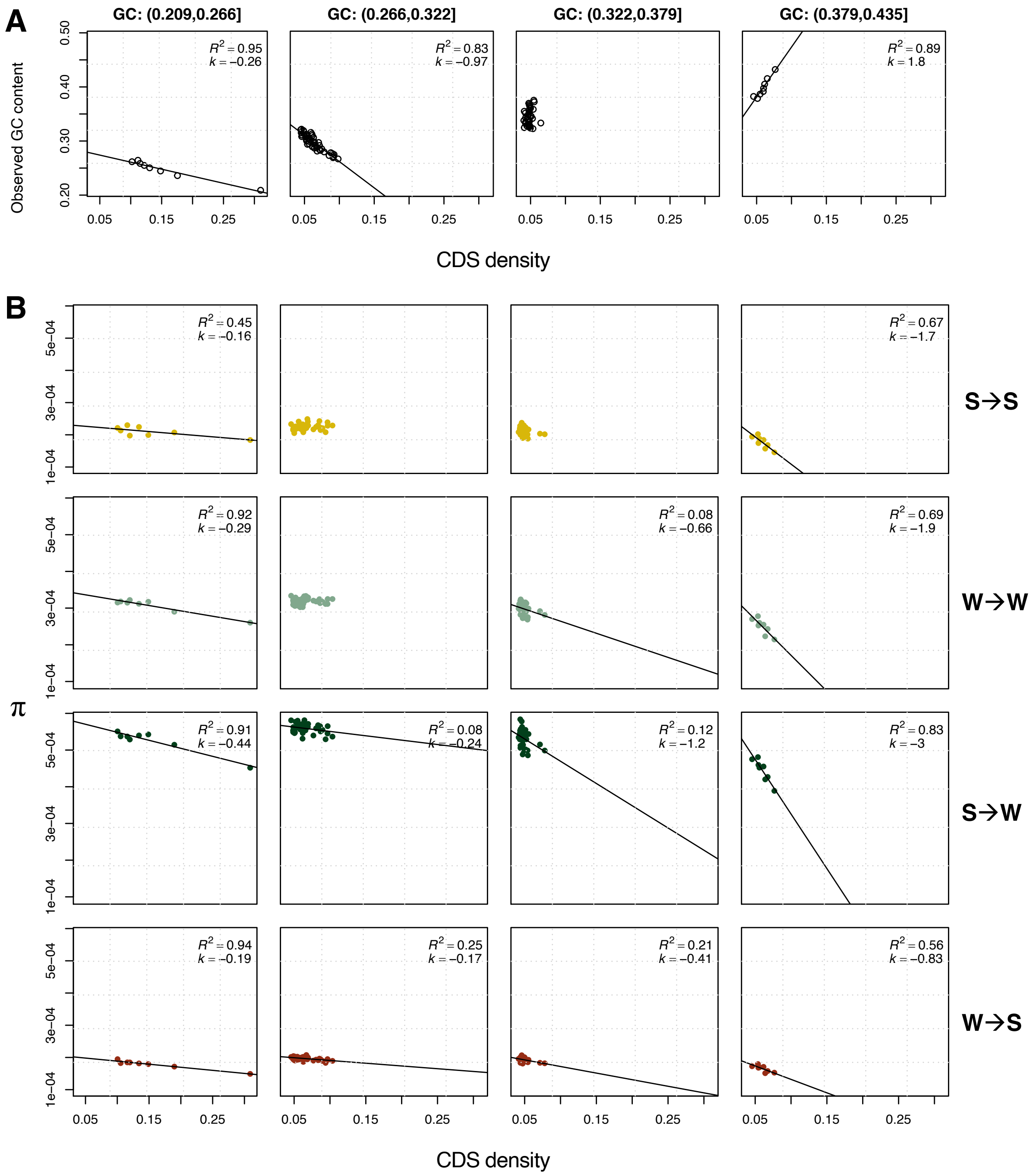


**Kaz-juv**

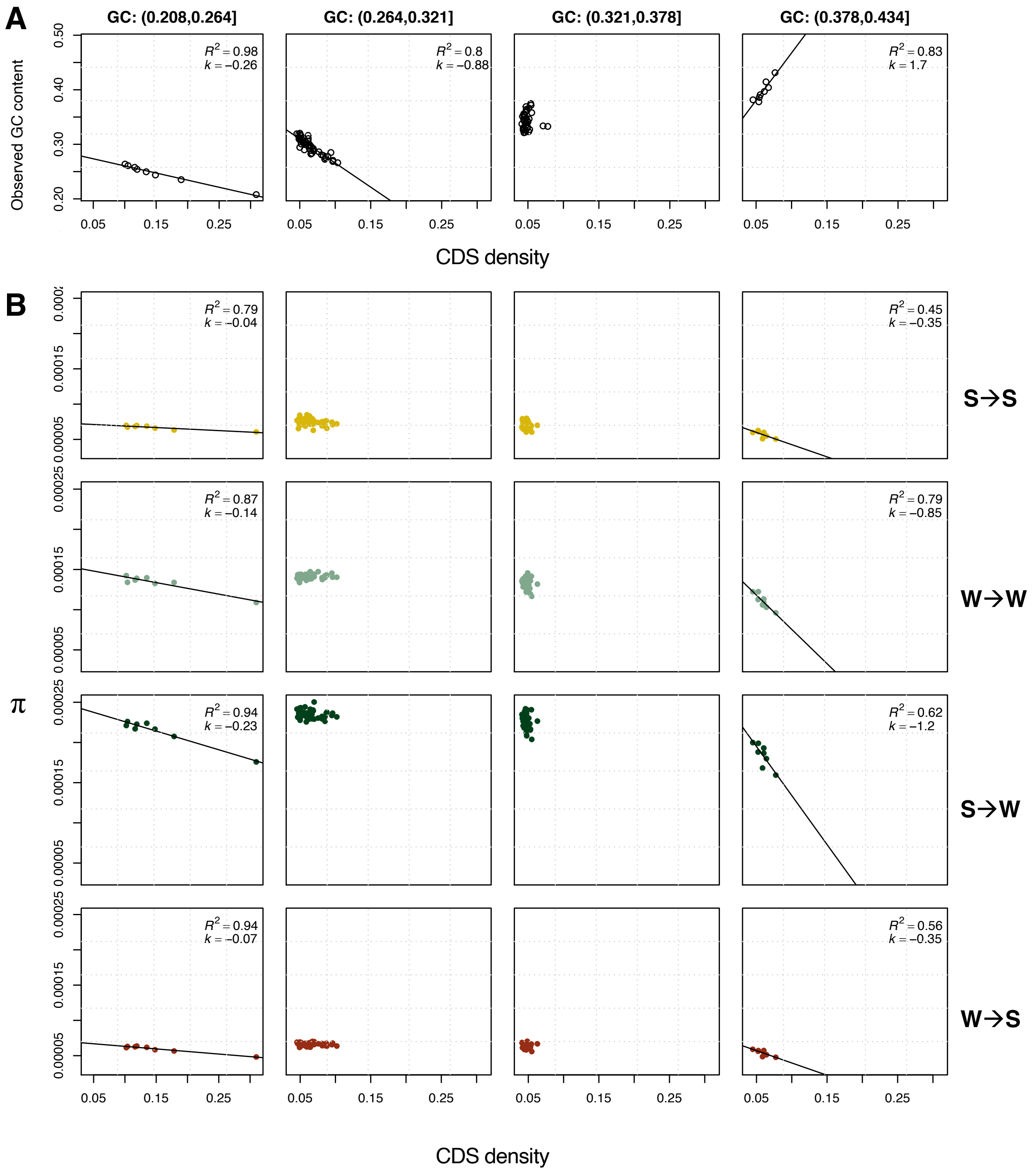
A
GC: (0.208,0.264]
GC: (0.264,0.321]
GC: (0.321,0.378]
GC: (0.378,0.434]
Observed GC content
$R^2$ = 0.98
$k$ = −0.26
$R^2$ = 0.8
$k$ = −0.88
$R^2$ = 0.83
$k$ = 1.7
CDS density
B
$R^2$ = 0.79
$k$ = −0.04
$R^2$ = 0.45
$k$ = −0.35
S→S
$R^2$ = 0.87
$k$ = −0.14
$R^2$ = 0.79
$k$ = −0.85
W→W
π
$R^2$ = 0.94
$k$ = −0.23
$R^2$ = 0.62
$k$ = −1.2
S→W
$R^2$ = 0.94
$k$ = −0.07
$R^2$ = 0.56
$k$ = −0.35
W→S
CDS density

**Ire-juv**

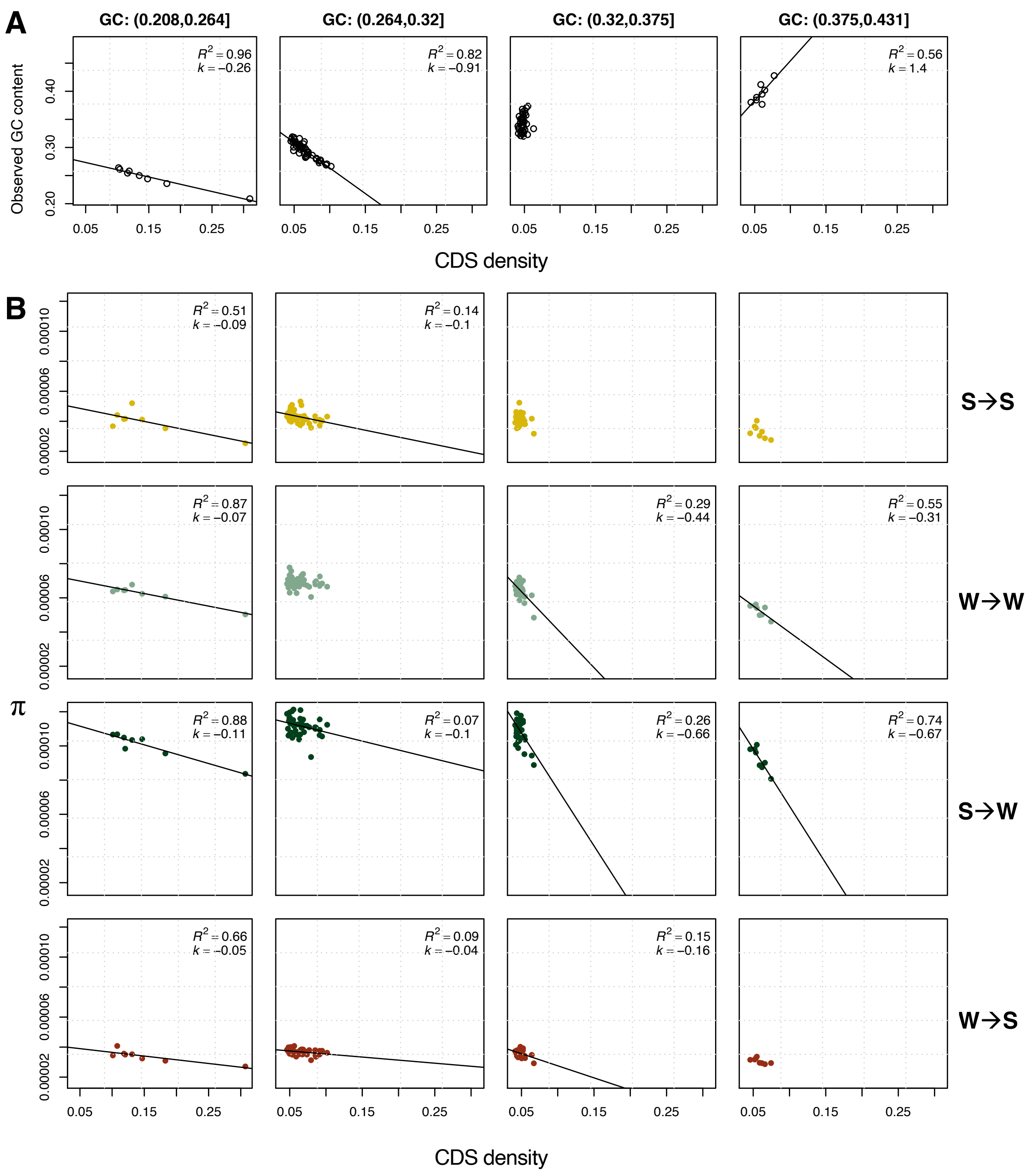
A
GC: (0.208,0.264]
GC: (0.264,0.32]
GC: (0.32,0.375]
GC: (0.375,0.431]
Observed GC content
R² = 0.96 k = −0.26
R² = 0.82 k = −0.91
R² = 0.56 k = 1.4
CDS density
B
S→S
W→W
S→W
W→S
π
R² = 0.51 k = −0.09
R² = 0.14 k = −0.1
R² = 0.87 k = −0.07
R² = 0.29 k = −0.44
R² = 0.55 k = −0.31
R² = 0.88 k = −0.11
R² = 0.07 k = −0.1
R² = 0.26 k = −0.66
R² = 0.74 k = −0.67
R² = 0.66 k = −0.05
R² = 0.09 k = −0.04
R² = 0.15 k = −0.16
CDS density

**Spa-rea**

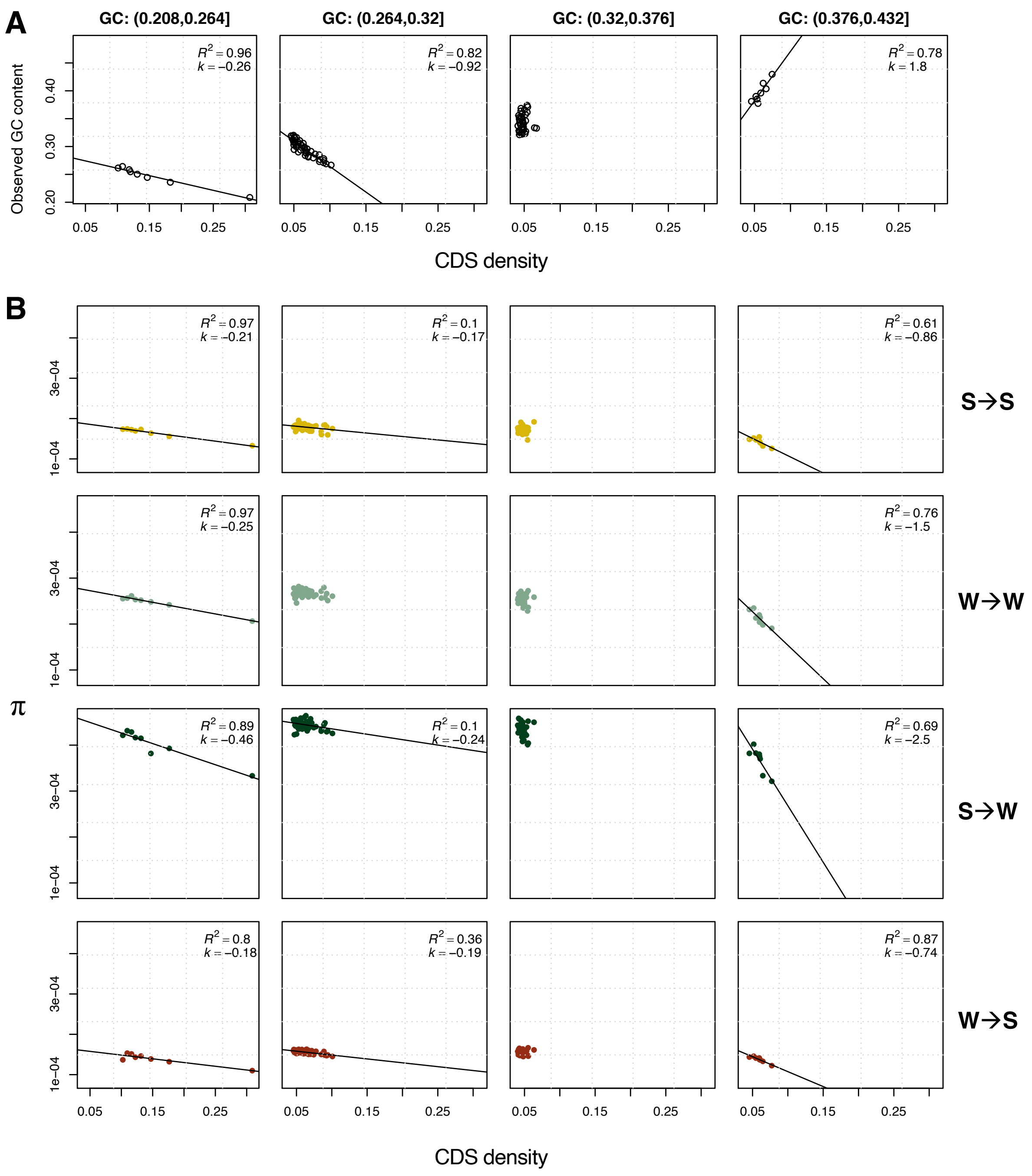

A
GC: (0.208,0.264]
GC: (0.264,0.32]
GC: (0.32,0.376]
GC: (0.376,0.432]
$R^2$ = 0.96
k = −0.26
$R^2$ = 0.82
k = −0.92
$R^2$ = 0.78
k = 1.8
Observed GC content
0.20
0.30
0.40
0.05
0.15
0.25
CDS density
B
$R^2$ = 0.97
k = −0.21
$R^2$ = 0.1
k = −0.17
$R^2$ = 0.61
k = −0.86
S→S
$R^2$ = 0.97
k = −0.25
$R^2$ = 0.76
k = −1.5
W→W
π
$R^2$ = 0.89
k = −0.46
$R^2$ = 0.1
k = −0.24
$R^2$ = 0.69
k = −2.5
S→W
$R^2$ = 0.8
k = −0.18
$R^2$ = 0.36
k = −0.19
$R^2$ = 0.87
k = −0.74
W→S
1e−04
3e−04
0.05
0.15
0.25
CDS density

#### Text S1: Mutation bias, CpG-sites and the impact of polarization scheme

##### Impact of polarization scheme on estimation of *B*

Previous analyses indicate that *B* might be overestimated when the polarization error rates are higher for S→W than W→S mutations (Glémin, et al. 2015). In our data, using the "strict" polarization scheme (see *Materials and Methods*), the W→S error rate was low (max 2.2% in Kaz-sin), but still higher than the S→W error rate (max 1.3%, also in Kaz-sin). This indicates that the M1 model for this data set underestimated *B* to some extent (Table S2). With a more liberal polarization approach, both W→S and S→W error rates increased to 7.6 % and 4.6% respectively (both for Kaz-sin). The W→S error was significantly higher in the liberal polarization ($p \approx 0.024$, paired *t*-test) but not the S→W error ($p > 0.05$, paired *t*-test). The M1* model was significantly better than the M1 model for all populations ($p < 0.05$, LRT). *B* from M1* was lower for the liberal polarization scheme for the *L. juvernica* populations: 0.71 (strict: 0.83) for Kaz-juv and 0.34 (0.57) for Ire-juv (M1*), but similar or slightly higher for the rest (Table S2). We conclude that the liberal polarization scheme provides qualitatively similar results but increases the polarization error rates.

##### Methodological considerations on the estimation of mutation bias

We found that λ varied from 3 – 4.1 among the *Leptidea* butterflies which is within the interval observed (2.1 - 4.5) across a wide range of organisms (Lynch 2007), but see also the extreme λ (11.69) observed in honey bees (Wallberg, et al. 2015) . Using the more liberal polarization approach (see *Materials and Methods*) lowered the λ of Ire-juv to 3 (from 3.5) and Kaz-juv to 3.5 (4.1) while the other populations remained unchanged. The strict polarization allows only private SNPs for the *L. juvernica* populations meaning that they consist of a mixture of sites that mutated after divergence or those derived alleles that were lost in either population. The advantage of the strict strategy is that sites are polarized with more confidence. Polarization errors has been shown to bias estimation of *B* (Glémin, et al. 2015) and could possibly bias estimation of λ as well. The disadvantages of the strict strategy are twofold but possibly less problematic than for the liberal polarization approach: i) fewer sites pass the filter ii) the DAF spectra are biased towards younger alleles. For i) this means a higher variance when estimating *B*, which did not appear problematic for the genome-wide values but probably reduces the power of inferences on the GC centiles. Point ii) could in theory be handled by the $r_i$ demographic parameters in the model which serves to correct for deviations in DAF spectra shared by all mutation categories (Eyre-Walker, et al. 2006; Muyle, et al. 2011). In practice, it is possible that the combination of a high λ and the signal of non-equilibrium demography imposed by polarization, pose difficulties for the model. This could potentially have been alleviated in Ire-juv, which shows a symptomatic excess of intermediate variants significative of a recent bottleneck (not shown). It is also possible that the GC content of the ancestral genome is underestimated for the *L. juvernica* populations. Compared to the reference Swe-sin population, *L. juvernica* show a widely divergent karyotype and consequently most likely changes in recombination landscapes. GC does not affect the estimate of *B* to a great extent, which is based on the shape of the DAF spectrum but underestimating the local GC content inflates the estimation of λ. However, this argument does not explain the lower λ observed for the *L. juvernica* populations when using the liberal polarization approach. To conclude, we cannot entirely rule out methodological issues but biological variation in mutation bias within *Leptidea* is possible.

**DNA methylation at CpG sites and mutation bias**

In general, we found a higher S→W mutation bias at ancestral CpG-prone sites indicating that these sites contribute more to the overall mutation bias (Table S2). To understand why this difference between trinucleotide classes exists, detailed knowledge about DNA methylation and mutation mechanisms in Lepidoptera will be needed (Bewick, et al. 2017; Jones, et al. 2018; Provataris, et al. 2018). Data from the cotton bollworm moth (*Helicoverpa armigera*), for example, show that only 0.9 % of CpG sites are methylated in this species and that CpG sites are underrepresented in methylated regions (Jones, et al. 2018). Jones et al. (2018) did not look specifically at germline tissue but did observe a lower CpG observed/expected ratio in methylated (0.738) compared to non-methylated (1.042), indicating that hypermutation of CpG sites could still be occurring in Lepidoptera.

**Impact of ancestral CpG-prone sites on estimation of *B***

In humans, CpG sites have a slightly higher level of gBGC when potential polarization errors are ignored (Glémin, et al. 2015). We therefore tested the impact of CpG sites on the estimation of *B* by splitting the non-exonic SNPs into two datasets: excluding ancestral CpG-prone sites and considering ancestral CpG-prone sites only. The GC content in the ancestral genome was ~0.21 and ~0.59 respectively when excluding ancestral CpG-prone sites or considering ancestral CpG-prone sites only. The difference in *B* between *L. juvernica* and the other species were amplified when excluding ancestral CpG-prone sites (Figure ST1A). For this data set, *L. reali* had a higher *B* than the *L. sinapis* populations, opposite to when all non-exonic sites were included (Table S2). When considering CpG-prone sites only, *B* estimates for both *L. juvernica* populations were lower than for the other SNP sets (Figure ST1B). For CpG-prone sites, the M1 model had a better fit than M1* for all populations except Ire-Juv (Table S2). In our analyses we found a higher *B* in most cases when using the polarization error correction model (M1*). This is an indication that CpG-prone sites are less likely to cause polarization errors in butterflies compared to species with more ubiquitous CpG-methylation such as humans.

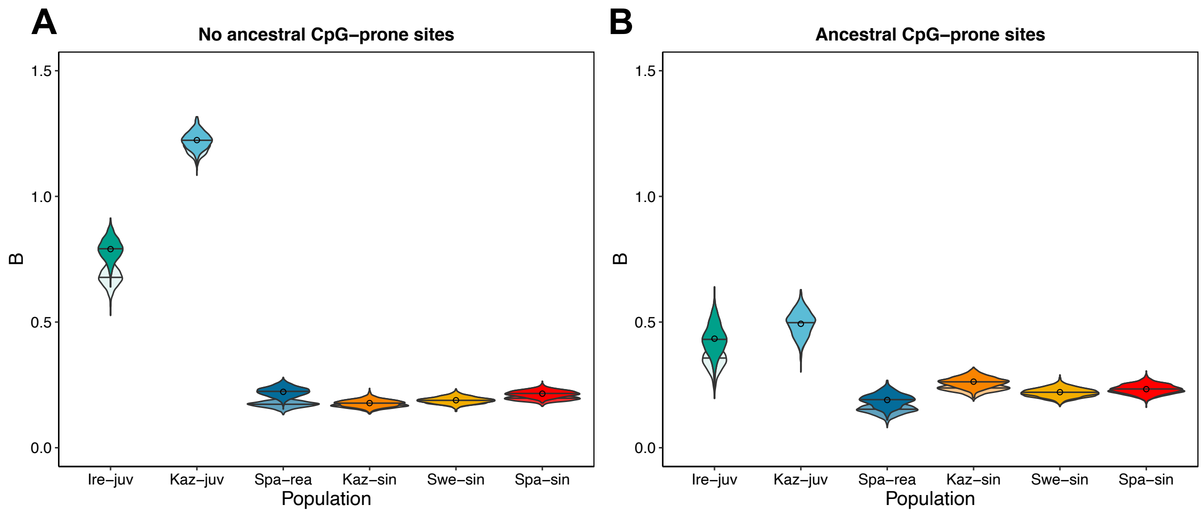


**Figure ST1. Estimates of *B* when A) excluding, and B) restricting analysis to ancestral CpG sites.** Circles represent point estimates from the original DAF spectra using model M1*. Bars are mean values of *B* for the 1,000 bootstrap replicates of sites.

### Supplementary references

Bewick AJ, Vogel KJ, Moore AJ, Schmitz RJ. 2017. Evolution of DNA methylation across insects. Molecular Biology and Evolution 34:654-665.

Eyre-Walker A, Woolfit M, Phelps T. 2006. The distribution of fitness effects of new deleterious amino acid mutations in humans. Genetics 173:891-900.

Glémin S, Arndt PF, Messer PW, Petrov D, Galtier N, Duret L. 2015. Quantification of GC-biased gene conversion in the human genome. Genome Research 25:1215-1228.

Jones CM, Lim KS, Chapman JW, Bass C. 2018. Genome-wide characterization of DNA methylation in an invasive lepidopteran pest, the cotton bollworm *Helicoverpa armigera*. G3: Genes, Genomes, Genetics 8:779–787.

Lynch M. 2007. The origins of genome architecture. Sunderland, MA: Sinauer Associates.

Muyle A, Serres-Giardi L, Ressayre A, Escobar J, Glémin S. 2011. GC-biased gene conversion and selection affect GC content in the *Oryza* genus (rice). Molecular Biology and Evolution 28:2695–2706.

Provataris P, Meusemann K, Niehuis O, Grath S, Misof B. 2018. Signatures of DNA methylation across insects suggest reduced DNA methylation levels in Holometabola. Genome Biology and Evolution 10:1185-1197.

Wallberg A, Glémin S, Webster MT. 2015. Extreme Recombination Frequencies Shape Genome Variation and Evolution in the Honeybee, Apis mellifera. PLoS Genetics 11:e1005189-e1005189.

Wickham H. 2016. Ggplot2: Elegant Graphics for Data Analysis. New York: Springer Verlag.
